# Benchmarking full-length ITS metabarcoding across Illumina 2x500, PacBio, and Oxford Nanopore sequencing using mock and soil communities

**DOI:** 10.64898/2026.05.20.726443

**Authors:** Leho Tedersoo, Marko Prous, Meirong Chen, Sten Anslan, Irja Saar, Benjamin Dubois, Vladimir Mikryukov

**Affiliations:** Mycology and Microbiology Center, Institute of Technology, University of Tartu, J. Liivi 2, 51409 Tartu, Estonia; Museum of Natural History, University of Tartu, Vanemuise 46, 51003 Tartu, Estonia; Ecology and Genetics Research Unit, PO Box 3000, 90014 University of Oulu, Finland; Institute of Ecology and Earth Sciences, University of Tartu, J. Liivi 2, 51409 Tartu, Estonia; Department of Biological and Environmental Science, University of Jyväskylä, Survontie 9, 40500 Jyväskylä, Finland; Biological Engineering Unit, Life Sciences Department, Walloon Agricultural Research Centre, 5030 Gembloux, Belgium

**Keywords:** fungi, high-throughput sequencing, Illumina MiSeq, PacBio Revio, sequencing bias, index switching

## Abstract

Metabarcoding is a powerful tool for biodiversity comparisons, where standard-size DNA barcodes (>500 bases) offer better taxonomic resolution than shorter ones. Still, the choice of sequencing platforms and bioinformatics pipelines may strongly affect inferred diversity due to various technical biases. We assessed the relative performance of Illumina MiSeq i100 (2x500 paired-end), PacBio Revio and Oxford Nanopore MinION sequencing and bioinformatics pipelines, using full-length ITS amplicon sequencing datasets from a 103-species mock community and 45 composite soil samples. Despite numerous low-quality reads, PacBio yielded the lowest overall error rate and highest number of taxa. Illumina revealed the highest proportion of chimeric and index-switched reads, along with a strong bias towards shorter amplicons. MinION data analysed using PRONAME and Minovar - a bioinformatics pipeline presented here - had the largest proportion of low-quality data, and rare taxa were lost during data filtering and read polishing steps. Although Minovar enabled amplicon sequence variant (ASV) level precision for common taxa, we recommend clustering ASVs into OTUs. For PacBio, standard filtering approaches outperformed the ASV approach because they retained rare taxa. For Illumina, a stringent ASV approach or removal of rare OTUs would limit artefacts. Across all platforms, excess PCR cycles promoted chimeric and low-quality reads and lost quantitativity in biodiversity assessments. With moderate differences in effect sizes, all analytical approaches supported the conclusion that sampling design determines how we see soil biodiversity responses to land use. For biodiversity surveys based on the full-length ITS metabarcoding, we recommend using PacBio sequencing with standard, non-ASV pipelines.

## 1. Introduction

Understanding biodiversity and its response to perturbations is one of the cornerstones in the disciplines of ecology, conservation biology and restoration biology (Tilman et al. 2014; Edwards & Cerullo 2024). Modern biodiversity assessment increasingly relies on molecular identification tools, particularly metabarcoding. Metabarcoding is a cost-efficient approach that relies on high-throughput sequencing (HTS) of a genetic marker from environmental DNA (eDNA) or bulk biological material, including soil, water, plant and animal samples. Metabarcoding is widely used to assess biodiversity patterns in microorganisms – bacteria, archaea, fungi and protists – as well as in macroorganisms, such as fish, arthropods and plants (Taberlet et al. 2012; Deiner et al. 2017). Metabarcoding techniques have been rapidly developing since the late 2000s, when HTS methods replaced DNA fingerprinting and Sanger sequencing-based methods (Taberlet et al. 2012).

The amplicon length is a critical issue in metabarcoding, because traditionally used, standard DNA barcodes, such as the internal transcribed spacer of the rRNA locus (ITS region) in fungi and certain protists and mitochondrial cytochrome oxidase I gene (COI) in animals are within 500-1000 bases and around 650 bases, respectively (Antil et al. 2023). However, short-read, second-generation HTS platforms, such as Illumina, Ion Torrent, DNBSeq and AVITI, enable sequencing of amplicons only up to 500-550 bases (Arslan et al. 2024; Kumar et al. 2024). Because these short-read methods cannot handle the standard barcodes, various minibarcodes - such as ITS1 and ITS2 subregions of the ITS, partial COI barcode and one or two neighbouring variable regions in the small and large rRNA subunit genes - became popular in eDNA metabarcoding analyses (Deiner et al. 2017). However, in all these cases, taxonomic resolution of the minibarcodes is substantially lower than that of the longer standard barcode (Jamy et al. 2020; Tedersoo et al. 2022; Varusk et al. 2025). To improve taxonomic resolution and consistency with specimen-based studies, it is therefore highly important to shift metabarcoding towards marker choices and data quality that better match standard DNA barcoding (Santoferrara et al. 2020; Tedersoo et al. 2022).

Recent third-generation, long-read HTS platforms, PacBio and Oxford Nanopore Technologies (ONT), have virtually no practical limit on amplicon size, given median raw read lengths of tens to hundreds of thousands of bases (Kumar et al. 2024; Zhang et al. 2024). Long-read methods have been considered less accurate, more expensive and more difficult to analyse compared with short-read methods (Tedersoo et al. 2021a). However, the 2022-2024 advances in hardware, sequencing chemistry and basecalling from ONT and PacBio have brought the data quality, quantity and cost of third-generation HTS data on par with Illumina sequencing (Zhang et al. 2025). On 20 October 2025, Illumina released 2x500 paired-end sequencing (Illumina 2025), which extends read length by ∼70% and, in theory, allows sequencing of amplicons exceeding 900 bases, including the standard ITS and COI barcodes.

Metabarcoding workflows have multiple quality-related issues, some inherent to amplification (i.e., PCR biases) and others specific to sequencing platforms and bioinformatics workflows. The main PCR errors include the incorporation of incorrect nucleotides, primer slippage and the formation of chimeric molecules, all of which are strongly influenced by the choice of polymerase and excessive amplification cycles (Santoferrara et al. 2020). For third-generation sequencing platforms, indels are a common issue, particularly at homopolymer positions (Tedersoo et al. 2021a; Zhang et al. 2025). By contrast, Illumina sequencing suffers from A-to-C transversions and errors associated with homopolymer ends and at specific motifs (Stoler & Nekrutenko 2021). Sequencing libraries of all HTS platforms use sample-specific indexed primers; hence, index-switching during library preparation or sequencing steps may be an important source of sample cross-contamination (Carlsen et al. 2012; Schnell et al. 2015; Loit et al. 2019).

Here, we used a technically replicated multi-species mock community and soil samples to evaluate the relative performance of Illumina 2x500, PacBio Revio, and ONT MinION in recovering the known biodiversity and producing technical artefacts. The main objective of this study was to validate alternative, cutting-edge sequencing methods and bioinformatics pipelines in metabarcoding analyses of fungi and protists based on the full-length ITS marker, which serves as the official barcode for fungi (Schoch et al. 2012) and one of the main barcodes for plants and multiple protist groups, such as dinoflagellates, ciliates, oomycetes and diatoms (Pawlowski et al. 2012; Antil et al. 2023). Additionally, we evaluated these sequencing methods to test an ecological hypothesis that the effect of land-use type on soil biodiversity patterns depends on sampling design. This question was motivated by unexpected results from a Europe-wide LUCAS soil survey showing that bacterial, fungal, animal and protist communities are more diverse in croplands compared with natural ecosystems, such as grasslands and forests (Labouyrie et al. 2023; Köninger et al. 2023; Aslani et al. 2024). Since these analyses were based on relatively small samples, i.e., five pooled soil cores from a 8-m^2^ area, it has raised doubts whether the observed pattern is scale-dependent and whether such a small sampling area is representative of the site in comparisons across land-use types (Froger et al. 2024; Chen et al. 2026).

## 2. Methods

### 2.1 Materials

To test the differences among DNA sequencing platforms, we used a mock community and 45 composite soil samples. The mock community consisted of a DNA mixture from 103 heterospecific fungal fruiting body specimens collected in various European countries in 2020-2024 (Table S1). DNA from these fruiting bodies was extracted from approximately 10 mg of dried material using an ammonium sulphate lysis protocol (Anslan & Tedersoo 2015).

Disregarding DNA concentration, 2 μL of DNA extract from each sample was pooled for high-throughput sequencing. For the reference sequences, the ITS region of each fruiting body specimen was separately amplified using the primers ITSOF (5’-acttggtcatttagaggaagt-3’) and LB-W (5’-cttttcatctttccctcacgg-3’), and Sanger-sequenced using the primers ITS5 (5’-ggaagtaaaagtcgtaacaagg-3’) and ITS4 at Macrogen Europe (Amsterdam, the Netherlands).

The soil samples were collected from five sites in each of the cropland, grassland and forest ecosystems (Table S2). At each of the 15 sites, three composite samples were collected using the protocols of the Global Soil Mycobiome consortium (40 subsamples, 5 cm diam. cores to 5 cm depth, 2500-m^2^ sampling area; Tedersoo et al. 2021b), SoilBON (9 subsamples, 5 cm diam. cores to 10 cm depth, 900-m^2^ area; Guerra et al. 2021) and European LUCAS soil survey (5 subsamples, 5 cm diam. cores to 20 cm depth, 8-m^2^ area; Orgiazzi et al. 2018). The central point of all three sampling designs was the same. The subsamples were pooled upon collection, thoroughly mixed, and about 100 grams of each composite sample was placed in a paper bag and dried in a heating cabinet with active airflow at 35 °C for 48 hours.

The dried composite soil samples were transferred to airtight plastic bags with zippers and further homogenised by hand rubbing. Around 1 g of finest particles was taken to an Eppendorf tube, supplemented by two tungsten balls, and subjected to further homogenisation by bead beating in Retsch MM400 homogeniser (Retsch GmbH, Haan, Germany) at 30 Hz for 5 min. Then, 0.20 g of soil dust was weighed for DNA extraction. DNA extraction from soil samples was performed using the Qiagen Soil Kit for the KingFisher robot (Qiagen, Carlsbad, CA, United States) according to the manufacturer’s instructions.

For mock community, composite soil samples, positive (Unispike artificial DNA molecule) and negative (double-distilled water) controls, amplification was performed using the primers ITS9MUNngs (5’-tacacaccgcccgtcg-3’) and ITS4ngsUni (5’-cctscscttantdatatgc-3’) under the PCR conditions outlined in Tedersoo et al. (2021b) with 30 cycles and 55 °C annealing temperature (hereafter, ‘PCR30’ treatment). All samples were additionally re-amplified using 35 PCR cycles (‘PCR35’) to represent a moderately excess-cycled technical variant. In all 96 amplicons, both primers were tagged with unique, symmetric 12-base indices. PCR products were checked on a 1.5% agarose gel and pooled according to band intensity, using 1-10 μL of PCR product. The PCR products were cleaned using FavorPrep GEL/PCR Purification Mini Kit (Favorgen Europe, Vienna, Austria) and subjected to platform-specific library preparation and sequencing.

### 2.2. Library preparation and sequencing

The PacBio SMRT cell library was prepared using the SMRTbell Prep kit v3.0, following the manufacturer’s instructions. The library was sequenced on one 25M SMRT cell along with another library, both occupying 50% of the SMRT cell capacity, on the Revio instrument using the SPRQ Polymerase kit and sequencing plate. Adaptive loading was used, and movie time was set to 24 hours. Circular consensus sequence (CCS) analysis for generating HiFi reads was performed on the instrument using the ICS SW v. 25.1.0.257715 software. PacBio library preparation and sequencing were performed at the Norwegian Sequencing Centre (divided service cost 1405 EUR).

For Illumina library preparation, we used the TruSeq DNA PCR-Free kit following the manufacturer’s instructions, except that the initial DNA fragmentation step was omitted. Sequencing was performed using the MiSeq i100 25M Reagent Kit (1000 cycles) on a MiSeq i100 instrument in the Illumina Solutions Centre Berlin. The flow-cell was shared with another library (50% cell space; divided service cost 2112 EUR).

For the ONT library preparation, the ONT Ligation Sequencing Kit V14 (SQK-LSK114) was used along with the NEBNext Companion Module for Oxford Nanopore Technologies Ligation Sequencing (E7180S). Sequencing was performed using the MinKNOW v25.05.14 (core 6.5.14) software on the MinION Mk1B instrument, using flow cell FLO-MIN114 version R10.4.1 for 24 hours (in-house cost 1071 EUR).

### 2.3. Bioinformatics

#### 2.3.1. Platform-specific bioinformatics analyses

PacBio HiFi reads were processed with the NextITS pipeline v1.1.0 (Mikryukov et al. 2026), executed in Nextflow v25.10.2 (Di Tommaso et al. 2017). Reads were demultiplexed using LIMA v2.12.0 (Pacific Biosciences) with “--min-score 85” option. The initial quality-filtering discarded sequences containing >4 ambiguous nucleotides, >0.01% expected errors, or homopolymer runs longer than 25 nucleotides. Primer trimming was performed with cutadapt v5.1 (Martin 2011), and reads lacking both primer sites in the correct orientation were removed.

Illumina paired-end reads were demultiplexed with cutadapt using 100% barcode overlap (12 bp), with both barcodes located within the first 30 bases of the forward and reverse reads, allowing up to 1 mismatch and no indels. Demultiplexed read pairs were merged using USEARCH v11.0.667 (Edgar 2010), requiring a minimum overlap identity of 80% and allowing up to 12 differences. After trimming the primers with cutadapt, merged reads were then processed with the NextITS pipeline as for the PacBio data.

ONT raw data (pod5) was basecalled with Dorado v1.2.0 using the highest accuracy model (dna_r10.4.1_e8.2_400bps_sup@v5.2.0) and enabling ‘--emit-moves’ option to include move tables in the output. The information in the move tables (per base dwell times) enables more accurate consensus sequence polishing with Dorado. Basecalled FASTQ reads were demultiplexed with cutadapt, allowing a minimum overlap of 11 bp and 1 mismatch. Subsequent steps differed between the two ONT pipelines.

Demultiplexed ONT data were imported into PRONAME v2.2.0 (Dubois et al, 2024), and only reads ranging from 450 to 3000 bp with a minimum Phred score of 15 were retained. Error correction was performed by first clustering reads at 98% identity using VSEARCH v2.22.1 (Rognes et al. 2016) and discarding singletons. Sequences were then polished with the polisher function of Dorado, using the latest and most accurate polishing model, taking into account move tables (dna_r10.4.1_e8.2_400bps_sup@v5.2.0_polish_rl_mv). The proname_refine script used at this step was slightly modified to deactivate chimera filtering, as this step was performed later in a common workflow for all datasets.

#### 2.3.2. Introducing the Minovar pipeline

We developed another ONT data analysis pipeline, Minovar (Prous & Mikryukov 2026), with a data-processing strategy that uses several clustering and polishing steps to determine sequence variants based on a minimum depth of 5 reads. Each demultiplexed input FASTQ file was processed separately to obtain consensus variant sequences. Cutadapt was used to recognize target amplicon reads based on the primer sequences (maximum error rate 0.15, minimum overlap 15, minimum output sequence length 400 bp). The reads were sorted with SeqKit2 v2.12.0 (Shen et al. 2024) by decreasing average quality score for subsequent VSEARCH clustering, which was performed first at 80% sequence similarity. The initial clustering similarity threshold was set low, because VSEARCH tends to oversplit clusters, creating many small clusters (possibly belonging to the same variant), which may be unreliable for consensus sequence creation. The resulting consensus sequences were then clustered at 93% sequence similarity to merge such oversplit clusters. For generating consensus sequences, 5-100 reads per cluster were used, as implemented in abPOA v1.5.6 (Gao et al. 2021). Clusters with <5 reads were excluded to avoid potentially error-prone consensus reads; likewise, from the clusters exceeding 100 reads, a random sample of 100 reads was taken for consensus creation to limit computation time and potential alignment inaccuracies at very high depth (Sievers et al, 2011; Wick, 2023). Identical consensus sequences and their corresponding reads were merged, followed by polishing the consensus sequences with Dorado, as described above for PRONAME. All reads in each cluster were mapped to the corresponding polished consensus sequences using minimap2 v2.30 (Li 2021), and single-nucleotide polymorphisms (SNPs) were detected with freebayes v1.3.10 (Garrison & Marth 2012). If high-quality SNPs (freebayes score >10) were detected, the reads were phased with devider v0.0.1 (nanopore-r10 preset; Shaw et al. 2025). Devider enables efficient detection of low-frequency variants. The original reads corresponding to the detected variants were then used to create consensus sequences separately and polished using abPOA and Dorado as described above (Figure 1).

**Figure 1.**
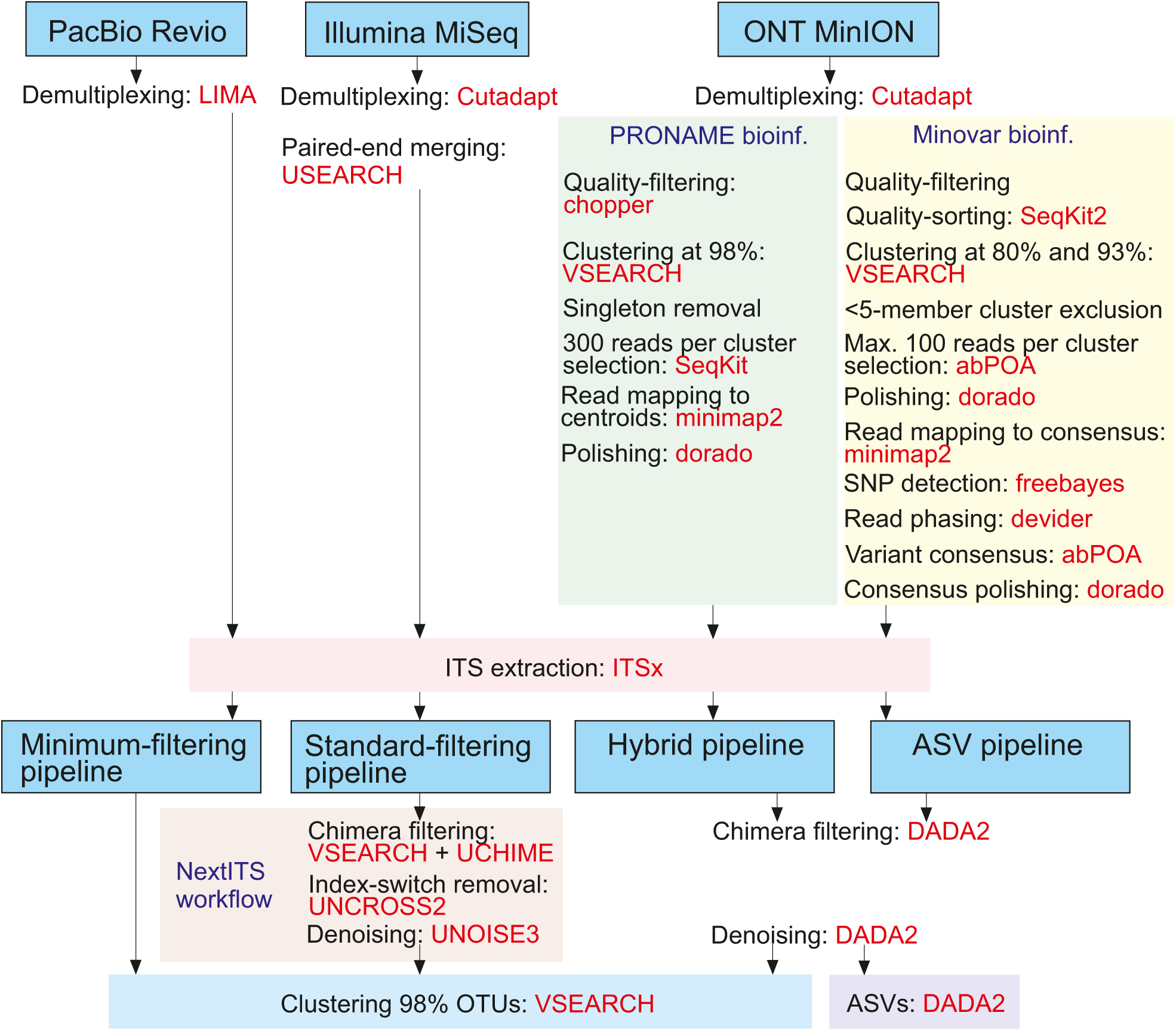
Bioinformatics workflow of the project. Blue typeface indicates analytical pipelines used; red typeface indicates specific programs used. Note that MinION data were not analysed using minimum-filtering and standard-filtering pipelines due to the initial low read quality.

#### 2.3.3. Common Bioinformatics for PacBio and Illumina data

General sequence handling (e.g., filtering, subsetting and format conversion) was performed using SeqKit2. Full-length ITS regions were extracted using ITSx v1.1.3 (Bengtsson-Palme et al. 2013) with the “all eukaryotes” setting and using the updated hidden Markov model (HMM) profile database (v2024, provided by R. Henrik Nilsson, University of Gothenburg, Sweden; profiles available at https://github.com/Mycology-Microbiology-Center/ITSx_HMMs). After ITSx extraction, representative ITS sequences were selected by retaining, for each group, the sequence with the highest mean Phred quality score computed with phredsort v1.3.0 (Mikryukov 2026): per-base Phred scores were first converted to error probabilities, the arithmetic mean error probability was calculated across positions, and this value was then transformed back to Phred scale to respect the logarithmic nature of Phred scores. The ITS-extracted data were further processed using four alternative bioinformatics pipelines.

To estimate the relative rates of chimera formation, low-quality reads and index-switching artefacts, we first used a ‘minimum-filtering’ approach that clustered ITS-extracted data with VSEARCH at 98% pairwise sequence similarity but without prior quality-filtering. As one of the baselines for taxonomic comparisons of PacBio and Illumina data, we used the ‘standard-filtering’ approach, with comprehensive quality-filtering as implemented in the NextITS workflow. Chimera removal was carried out in two stages using VSEARCH v2.30.0: (i) *de novo* detection with the UCHIME algorithm (Edgar et al. 2011) using a maximum chimera score of 0.6, followed by (ii) reference-based validation (uchime_ref) against the EUKARYOME v2.0 database (Tedersoo et al. 2024); sequences flagged as chimeras in either step were excluded. Within each sample, sequences differing only in homopolymer length were consolidated, and the most abundant variant was retained as the representative sequence. Within sequencing runs, index-switching (tag-jump) artefacts were filtered with the UNCROSS2 algorithm (Edgar, 2018) using ‘f = 0.01’. Before clustering, ITS sequences shorter than 150 bp or with >0.6 expected errors per 100 bp were removed. The remaining reads were denoised with the UNOISE3 algorithm (Edgar, 2016) with parameters ‘alpha = 6’ and ‘minsize = 1’, and subsequently clustered with VSEARCH at 98% pairwise sequence similarity (Figure 1).

As another baseline for among-platform comparisons, we used the DADA2 approach (Callahan et al. 2016) to infer amplicon sequence variants (ASVs) from Illumina and PacBio reads, producing representative sequences that are comparable to the polished ONT consensus sequences. For both platforms, reads were analysed in single-end mode following the ITS amplicon workflow of DADA2 v1.36.0. Error rates were estimated using the ‘loessErrfun’ function, and ASV inference was performed independently for each sample (‘pool = FALSE’). DADA2 algorithm options were set using ‘setDadaOpt’, including a band size of 16 for banded Needleman-Wunsch alignment, singleton detection disabled and denoising stringency parameters ‘OMEGA_A=1e-40’, ‘OMEGA_C=1e-40’ and ‘OMEGA_P=1e-04’.

To further compare the ‘standard-filtering’ and DADA2 denoising pipelines, we used a hybrid approach in which ASVs were clustered into 98% sequence-similarity OTUs as described above (Figure 1). This additional clustering step collapses the substantial intraindividual and intragenomic ITS variability typical of fungi and reduces richness inflation that can arise when each ITS haplotype is treated as an independent unit, while retaining the error-aware denoising benefits of DADA2 (Kauserud 2023).

For taxonomy annotation, representative sequences of each OTU or ASV from all four sequencing platforms and four bioinformatics treatments were subjected to BLASTn (Camacho et al. 2009) against EUKARYOME v2.0 as a reference database. Taxon names were assigned from the best hit, using sequence similarity and e-value thresholds (Tedersoo et al. 2021b). For the mock community, we also used BLASTn against the Sanger-sequenced reference of all specimens and representative sequences of all soil OTUs separately. In addition, for both mock and soil communities, we performed BLASTn searches of the ITS1 region and the ITS2 subregion (obtained via ITSx) separately to identify chimeric OTUs.

### 2.4. Data curation

For the mock community, OTUs were considered as successfully recovered when their representative sequences had up to 2 mismatches to the Sanger-sequenced reference. OTUs were considered chimeric when the ITS1 and ITS2 subregions matched to a different reference at >99% sequence similarity or the match coverage was <90%. Non-chimeric OTUs with >2 mismatches to the Sanger-sequence reference were considered of low quality or index-switched. Index-switched OTUs were defined as OTUs with <99% sequence similarity to the Sanger reference but with a stronger match to any member of the mock community or positive control. A few outlying OTUs, present in most mock community datasets but absent from the soil community, were considered field contaminants of the fruiting body samples. These were excluded from subsequent analyses.

For the soil community, data quality was assessed in 7 steps using OTU data and BLASTn output. First, OTUs with >97.0% match coverage and >99.0% sequence similarity to any reference sequence were assigned as high quality. Second, OTUs with a Phred score <30.0 were assigned as low-quality (except in ONT data). Third, OTUs with gap openings exceeding substitutions by >2 for all 3 best-matching sequences were assigned to low quality. Fourth, OTUs with genus-level to kingdom-level taxonomic mismatches in ITS1 and ITS2 were considered chimeric if their e-value multiplication was <e50 (i.e., identification of at least one subregion was reliable enough). Fifth, OTUs with <95.0% match coverage but >99.0% sequence similarity to any reference sequence were assigned as chimeric. Sixth, OTUs with >99.0% sequence similarity in ITS1 but <96% sequence similarity in ITS2, or vice versa, were considered chimeric. Seventh, OTUs for which the average of ITS1 and ITS2 sequence similarities exceeded the full-ITS sequence similarity by >1 percentage point were assigned as chimeric. All other OTUs were considered potentially high-quality. For each platform-bioinformatics data combination, the remaining index-switching rate was estimated using the default UNCROSS scores and the presence of positive-control reads in biological samples.

### 2.5. Data analysis

To evaluate the relative performance of sequencing platforms and bioinformatics pipelines on data quality, we calculated the following variables for each sample: the number of chimeric sequences and OTUs, the number of low-quality sequences and OTUs, the number of index-switched sequences and OTUs and the number of OTUs matching the reference dataset at 100.0% identity and coverage. Across all samples, we also considered the lengths of representative sequences. To test ecological hypotheses, we calculated fungal OTU and genus richness, the fungal Shannon index and eukaryote OTU and genus richness. To account for differences in sequencing depth, richness values were converted to residuals based on log-log regressions against sequencing depth (Tedersoo et al. 2022). Relative abundances of chimeric, low-quality and tag-switched reads were log-ratio-transformed relative to the numbers of high-quality sequences. Logarithmic and log-ratio transformations were used to improve the distribution of residuals and reduce internal dependencies in proportional data, respectively.

For each transformed technical and ecological variable, we built generalised linear mixed models (GLMMs) with a Gaussian error structure, where sequencing platform (df=3), bioinformatics pipeline (df=3), PCR cycle number (df=1), sampling design (df=2) and land-use type (df=2) served as fixed factors and study site (df=15) served as a random factor. Two-way interactions were also fitted, with a focus on the interaction term between sampling design and land-use type.

> *Data quality or richness* ∼ *PCR cycle number + Sampling design + Sequencing platform + Bioinformatics pipeline + land-use type + Sampling design * land-use type + (1*∣*Study site)*

In the analyses of fungal Shannon index, log-transformed sequencing depth was used as a covariate.

> *Fungal Shannon index ∼ PCR cycle number + Sampling design + Sequencing platform + Bioinformatics pipeline + land-use type + Sampling design * land-use type + sequencing depth + (1*∣*Study site)*

Besides these overall models, we calculated local models for each data subset as divided by combinations of sequencing platform, bioinformatics pipelines and PCR cycles. To test the effects of technical predictors on consistency of their ecological explanatory power, we used determination coefficients for sampling design, land-use type and their interaction from these local models as dependent variables.

> *Data quality indicator ∼ Sampling design * land-use type + (1*∣*Study site)*

Pearson correlation tests were used for pairwise relationships between read length, richness, OTU/ASV abundance and artefact proportions. The analyses were performed in R v4.3.1 (R Core Team 2023) using the stats v4.3 and lme4 v1.1-35.1 (Bates et al. 2015) packages

## 3. Results

### 3.1. Overall statistics

PacBio (half of a SMRT cell), Illumina (half of a run) and ONT (full, 24-hour run) recovered 6,868,925, 11,582,494 and 6,433,620 raw reads, respectively. For PacBio, the average polymerase read length was 77,920 bases, enabling an average 84-fold CCS and an expected accuracy of >99.9%. For the polished ONT reads, the median accuracy was estimated at 98.0-98.5% based on the mock community, whereas 1.3-1.6% of the reads were identical to Sanger sequences (i.e., error-free). The demultiplexing rate was highest in Illumina (81.1%), followed by PacBio (59.5%). Despite allowing 1 bp of index mismatch (no mismatches allowed for Illumina and PacBio data), the demultiplexing yield was lowest for ONT data at 39.3%.

Primers were successfully detected in 96.3% of demultiplexed PacBio reads and 92.5% of Illumina reads but only 76.6% of ONT reads. Unexpected or artefactual index combinations occurred in 1.2% of PacBio reads, 2.0% of ONT reads and 5.7% of Illumina reads. Altogether 4,015,687 (58.5%) PacBio sequences, 9,298,978 (80.3%) Illumina sequences and 1,687,141 (26.2%) ONT sequences passed these initial quality-filtering steps and were used in further analyses. Of the retained reads, 1.4% of Illumina reads, 0.5% of PacBio reads and 0.4% of ONT reads were predicted to be index-switched based on UNCROSS scores and the distribution of positive-control OTUs in biological samples.

There were some differences in read distribution across samples. While the sequencing depth among PacBio samples varied 9.7-fold (from 8960 to 87601 reads), the differences in Illumina and ONT were 14.4-fold and 20.2-fold, respectively. Similarly, the coefficient of variation in sequencing depth per sample was lower for PacBio (CV=0.44) than Illumina (CV=0.49) and ONT (CV=0.61).

### 3.2. Mock community

The mock community provided the controlled part of the benchmark, allowing us to separate recovery of known taxa from artefacts introduced by amplification, sequencing and bioinformatics. Of 103 specimens in the mock community, 97 were successfully recovered using various HTS technologies, with differences among sequencing platforms, PCR cycle numbers and bioinformatics workflows (Table S1). In addition to the input taxa, five species of yeast fungi and apicomplexans were consistently found and removed from subsequent analyses as contaminants. These organisms probably colonised one of the fruiting bodies, since they were not perfectly matched to any taxa detected in the soil samples.

In the standard-filtering and minimum-filtering approaches, PacBio revealed 93-96 input taxa, while Illumina recovered 89-90 (Table S1). Notably, Illumina sequenced five additional, medium-abundance taxa, but their representative sequences had 3-6 mismatches to the mock standard. In the ASV-based approach, PacBio and Illumina recovered 84-92 and 82-90 input taxa, respectively. Minovar and PRONAME recovered 75-84 and 81-82 taxa, respectively, in the ONT platform. Nonetheless, the total number of ONT high-quality ASVs reached 93 across PCR and bioinformatic treatments. Most of these were low-abundance taxa, indicating inconsistencies in how the ONT bioinformatics pipelines handle rare species.

In the minimum-filtering pipeline, the number of high-quality OTUs overestimated the recovered number of input taxa by 2.7-fold to 2.8-fold in PacBio and 1.6-fold to 1.7-fold in Illumina. In the PacBio platform, standard-filtering, ASV and hybrid approaches reduced these overestimates to 3.2-12.3%, 26.1-26.2% and 1.1-1.2%, respectively. The respective values for Illumina were similar, 9.9-15.5%, 23.3-29.3% and 0.0-2.5%. For the ONT data, the ASV and hybrid approaches overestimated the target taxon richness by 20.0-33.3% and 1.3-3.6%, respectively, in the Minovar pipeline, and by 92.6-141.5% and 1.2-2.4% in the PRONAME pipeline. These estimates suggest that for the ONT data, Minovar offers nearly ASV-level resolution in the complex mock community. We hypothesize that the 20-35% richness overestimation in the ASV pipeline across all platforms reflects intraindividual and/or intragenomic ITS variation (Smith et al. 2007).

PCR cycle numbers had variable, inconclusive effects on the recovery of input taxa. In standard- and minimum-filtering approaches, the PCR35 treatment (35 cycles) recovered 1-4 more taxa compared with PCR30 (30 cycles). However, the ASV approach revealed 8 fewer taxa for both PacBio and Illumina at PCR35. In ONT-Minovar and ONT-PRONAME, respectively, PCR35 yielded 9 and 1 additional taxa. Similar trends were evident in the hybrid approach (Table S1). These differences in recovery were nearly always attributable to the relatively low-abundance species. The taxa that were not recovered were therefore most likely lost through a combination of unequal template representation, PCR bias, stochastic loss of rare templates and downstream filtering. In particular, DADA2 denoising and ONT consensus generation are expected to remove uncommon variants, whereas filtering for chimeras and low-quality reads may further remove rare taxa that lack database matches or have low-quality scores.

Unlike other platforms, there was a negative correlation between read length and OTU abundance in Illumina data, which was more pronounced at PCR35 (ASV approach: R=-0.531; P<0.001) than at PCR30 (R=-0.387; P<0.001; Figure S1). At PCR30, Illumina sequence abundances were moderately to strongly correlated with abundances in PacBio (R=0.609, P<0.001), ONT-PRONAME (R=0.733, P<0.001) and ONT-Minovar (R=0.726, P<0.001), but these relationships weakened at PCR35 (R range 0.130 to 0.341). Meanwhile, correlation coefficients in sequence abundance between PacBio and ONT pipelines ranged from 0.935 to 0.968 in the ASV approach. PCR30 and PCR35 exhibited strong, overall positive correlations across all platforms (Illumina: R=0.882; PacBio: R=0.989; ONT-PRONAME: R=0.961; ONT-Minovar: R=0.963).

The rate of artefact accumulation varied greatly among sequencing platforms, bioinformatics pipelines and PCR treatments. In the minimum-filtered data, 1.23% of PacBio reads (17.7% OTUs) and 3.38% of Illumina reads (48.4% OTUs) were considered chimeric at PCR30 (Figure 2). Standard-filtering reduced the chimerism rate to 0.05% (3.2% OTUs) and 0.09% (2.9% OTUs), respectively. In the ASV approach, the chimerism rates were 0.00%, 0.51% (21.5% OTUs), 0.05% (0.86% OTUs), and 0.00% for PacBio, Illumina, ONT-PRONAME, and ONT-Minovar, respectively. At minimum filtering, relative chimera abundance increased 7-fold to 14-fold in the PCR35 treatment in Illumina and PacBio, reaching 17.0-23.7% of all reads. These differences between PCR30 and PCR35 were similar in the ASV pipeline for Illumina and PacBio. In ONT workflows, the proportion of chimeric sequences increased from 0.0-0.9% in PCR30 to 7.2-7.5% in PCR35.

**Figure 2.**
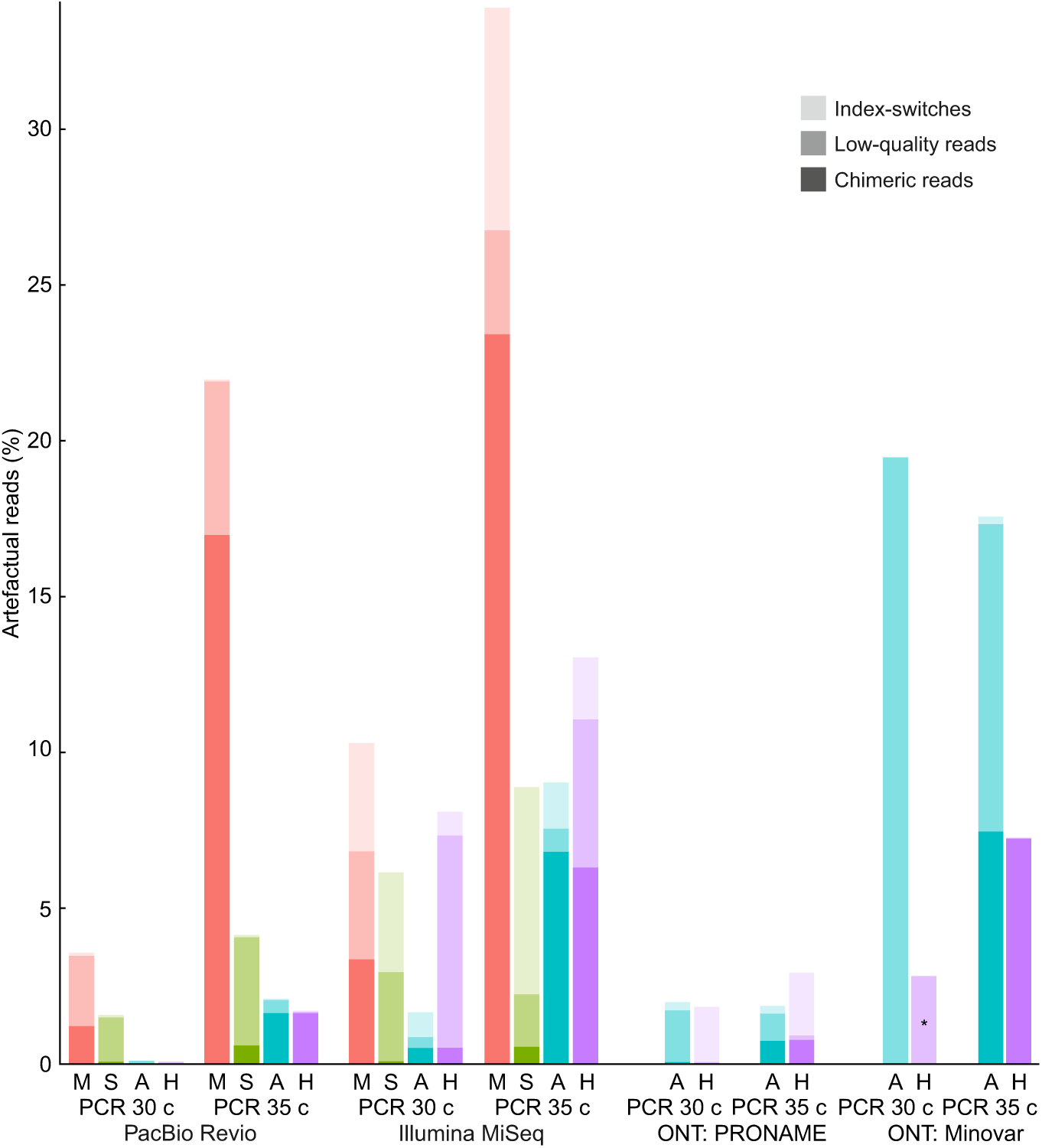
Proportions of artefactual reads as deduced from artefactual OTUs, depending on sequencing platform, bioinformatics pipelines and PCR cycle number. Abbreviations: M, minimum-filtering; S, standard-filtering; A, ASV; and H, hybrid approach. An asterisk indicates a sole effect of a single high-abundance OTU.

Based on low-quality OTUs, we calculated the proportions of low-quality reads. Low-quality reads accounted for 2.2% to 4.8% of all sequences across the PacBio and Illumina platforms, with no clear differences between platforms or PCR cycles (Figure 2). These figures were reduced to 1.4-3.5% in the standard OTU pipeline and to 0.1-0.7% in the ASV pipeline. Low-quality reads accounted for 0.9-1.7% of sequences in ONT-PRONAME and 9.9-19.3% in ONT-Minovar, indicating shortcomings in consensus building with unfiltered reads. Unlike in other platforms, the hybrid approach virtually eliminated low-quality reads and OTUs in the ONT data.

Index switching was a major issue for Illumina compared with other platforms (Figure 2). In the minimum filtering pipeline, index switches accounted for 3.47-7.14% of reads (18.4-36.6% OTUs) in Illumina sequencing and 0.07-0.10% reads (0.33-2.27% OTUs) in PacBio sequencing, with no effect of PCR treatments. In the standard-filtering pipeline, these values dropped by only 20%, but the proportion of index-switched OTUs increased by 1.9-fold to 4.3-fold due to the more efficient removal of other artefacts. In the ASV pipeline, index-switching contributed 0.00-0.05% (0.0-2.5% OTUs), 0.77-1.99% (13.6-15.6%), 0.26-0.27% (5.4-5.6%) and 0.00-0.22% (0.0-2.1%) of the data in the PacBio, Illumina, ONT-PRONAME and ONT-Minovar approaches, respectively. Similar values were evident in the hybrid approach.

Although there was no ground truth for taxon relative abundances, we calculated the CV of individual taxon abundances for comparative purposes. In the ASV and hybrid pipelines, the CV of OTU abundances was distinctly lower for Illumina (median, 1.48; range, 1.27, 1.74) than ONT (1.92; 1.82, 2.75) and PacBio (2.44; 1.96, 2.93). In PCR35, the CV values were 6.9%-50.8% greater than in PCR30, indicating reduced evenness with surplus cycles.

### 3.3. Soil microbiome

#### 3.3.1. General results

The soil data extended the mock community benchmark to a complex field setting, where exact taxon recovery are unknown, but the consequences for biodiversity estimates and ecological contrasts can be evaluated. The PacBio standard-filtering approach recovered on average 2621±880 eukaryote OTUs assigned to 453±115 genera, including 298±87 fungal genera. Sequencing platforms and bioinformatics pipelines exhibited marked differences in the recovery of soil biodiversity. Across all treatments, PacBio revealed an average of 1691±665 (SD), 1233±510, 384±159 and 344±137 fungal OTUs per sample in the minimum-filtering, standard-filtering, ASV and hybrid approaches, respectively. Using the ASV approach, Illumina recovered on average 99.3% more ASVs, whereas ONT-PRONAME and ONT-Minovar recovered 6.0% and 60.7% fewer ASVs, respectively, than PacBio (Figure 3).

**Figure 3.**
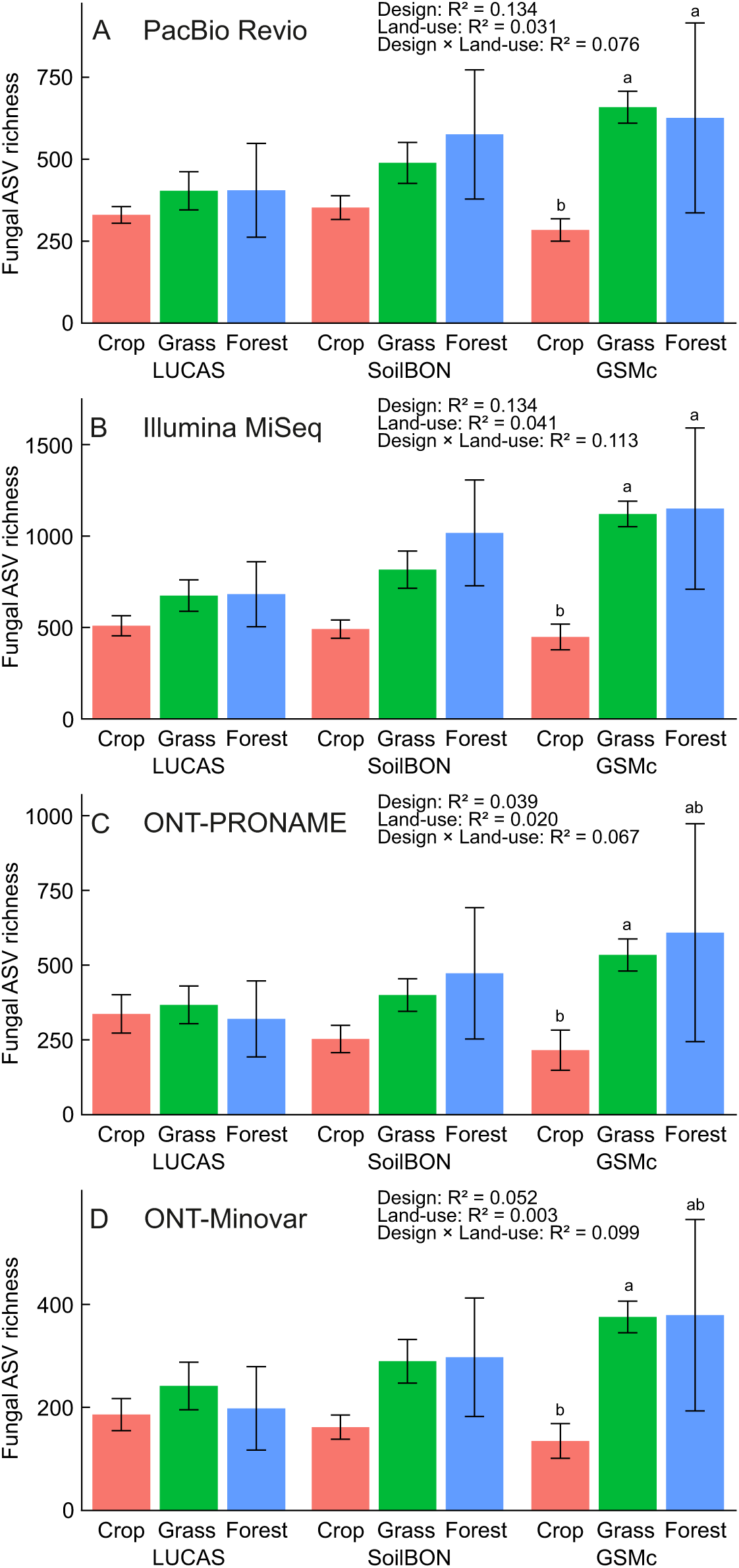
Relative performance of sequencing platforms in determining the effect of sampling design, land-use and their interaction on fungal ASV richness at PCR30. A) PacBio; B) Illumina; C) ONT-PRONAME; and D) ONT-Minovar approaches (n=60 for each bar).

Across all sequencing platforms and bioinformatics pipelines, fungi were the dominant group (average, 55.0% of OTUs), followed by ciliates (11.8%) and animals (7.7%). However, the proportions of most kingdom- and phylum-level groups varied mainly by the sequencing platform, with PacBio differing most from ONT and Illumina platforms (Figure 4). Compared with the average, PacBio revealed 34.4% lower richness proportion of apicomplexans, 25.8% lower proportion of ciliates, 17.8% lower proportion of fungi (including 27.9% less ascomycetes, 21.2% less basidiomycetes and 30.4% less glomeromycetes) but 75.0% more amoebozoans and 88.0% more cercozoans. Illumina recovered a richness proportion of early-derived fungal lineages 25.5% higher than in PacBio and 61.1% higher than in ONT, and a richness proportion of euglenids 55.7% higher than in PacBio and 200.0% higher than in ONT. Across all platforms, the hybrid approach revealed 81.5%, 69.4% and 31.3% greater proportional richness of euglenids, amoebozoans and early-diverging fungal lineages, respectively, compared with the ASV pipeline. The differences in hybrid and ASV approaches indicate differences in intraspecific diversity among groups of organisms.

**Figure 4.**
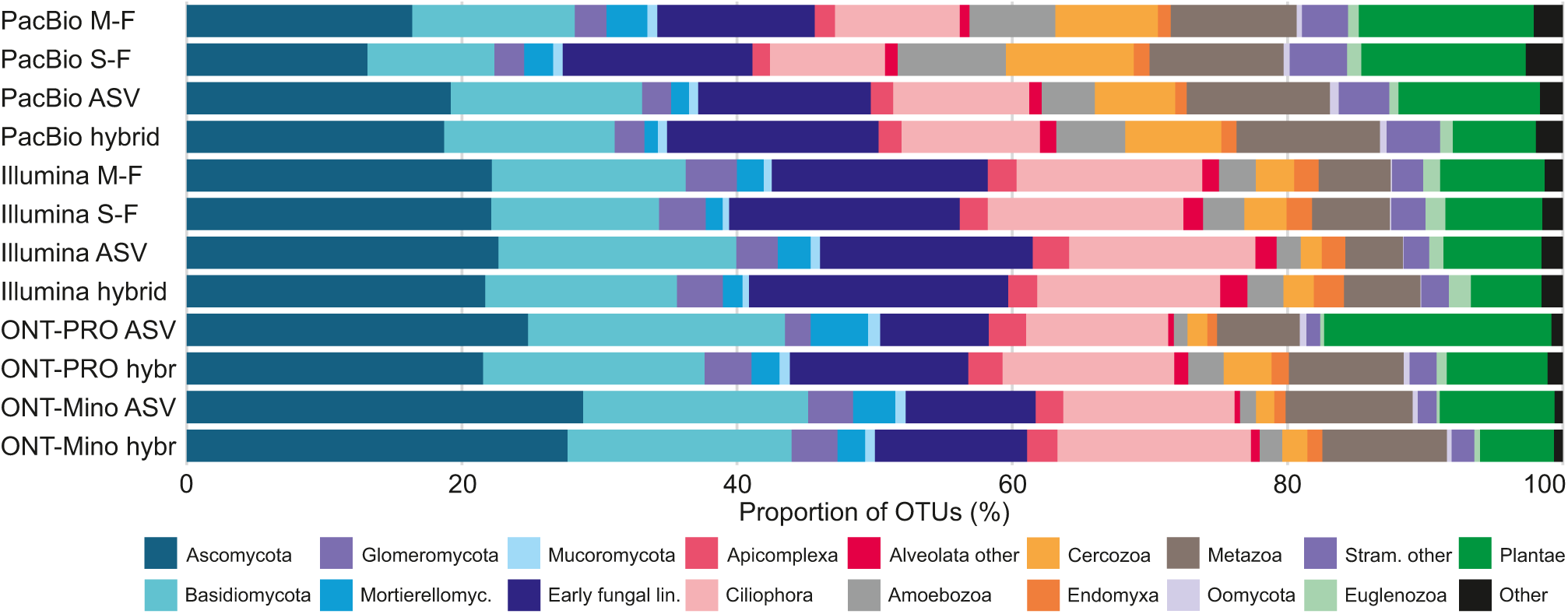
Proportions of major eukaryote groups and fungal phyla recovered by various sequencing platforms and bioinformatics approaches. Abbreviations: M-F, minimum-filtering; S-F, standard-filtering; ONT-PRO, Oxford Nanopore PRONAME; ONT-Mino, Oxford Nanopore Minovar.

At the OTU level, sequencing platforms and bioinformatics approaches agreed that *Aegopodium podagraria* is the most common taxon, but the ranking of other taxa varied greatly (Table S3). For the top 50 OTUs, PacBio data were more strongly correlated with ONT-Minovar (R=0.699) and ONT-PRONAME (R=0.601) than with Illumina (R=0.467) in the ASV treatment. Alarmingly, the standard-filtered PacBio and Illumina data had a poor correlation (R=0.206). Much lower read proportions of certain vascular plant OTUs (e.g., *Viola tricolor*, *Alopecurus pratensis*, *Achillea millefolium* and *Tilia cordata*) in the Illumina data mainly corresponded to these differences. Within both Illumina and PacBio datasets, the OTU rankings were well correlated across standard-filtering, minimum-filtering and hybrid datasets (R range 0.783 to 0.965), but only moderately correlated with the ASV datasets (R range 0.608 to 0.637). Here, the key difference is the presence of multiple ASVs in certain plant species (e.g., *Brassica rapa* and *Taraxacum officinale*) and lumping several biological fungal species, represented by distinct ASVs, into OTUs (e.g., *Fusarium avenaceum* and *Gibellulopsis nigrescens*). Furthermore, organisms that ranked lower or were absent in the Illumina sequencing had amplicons longer than average.

OTU richness estimates were moderately to strongly correlated among sequencing platforms (fungi at PCR30: R range 0.612 to 0.818; eukaryotes at PCR30: R range 0.767 to 0.916) and bioinformatics techniques (fungi at PCR30: R range 0.847 to 0.987; eukaryotes at PCR30: R range 0.789 to 0.998), with no difference among these technical approaches (Figure S2). Richness estimates based on PCR30 and PCR35 were less strongly correlated (fungi: R range 0.340 to 0.569; eukaryotes: R range 0.603 to 0.769; Figure S3).

#### 3.3.2. Compromised sequences and OTUs

Artefact profiles in the soil data broadly reflected the mock-community results, but the absolute rates were lower because artefact recognition in complex communities was necessarily more conservative. Across sequencing platforms and bioinformatics pipelines, chimeric sequences contributed to 1.53% of all reads (Figure 5A). The proportion of chimeric sequences was 3.0-fold higher at PCR35 than PCR30 (Overall mixed model: R^2^=0.039; P<0.001; Table S4), 2.2-fold higher in Illumina and 1.4-fold higher in ONT than PacBio (R^2^=0.140; P<0.001). Compared with the standard-filtering approach, the proportions of chimeric sequences were 8.3-fold, 2.8-fold and 2.9-fold higher in the minimum-filtering, hybrid and ASV approaches, respectively (R^2^=0.227; P<0.001). A significant interaction term indicated that more chimeric molecules were sequenced with Illumina compared with PacBio and ONT at high PCR cycle numbers (R^2^=0.019; P<0.001).

**Figure 5.**
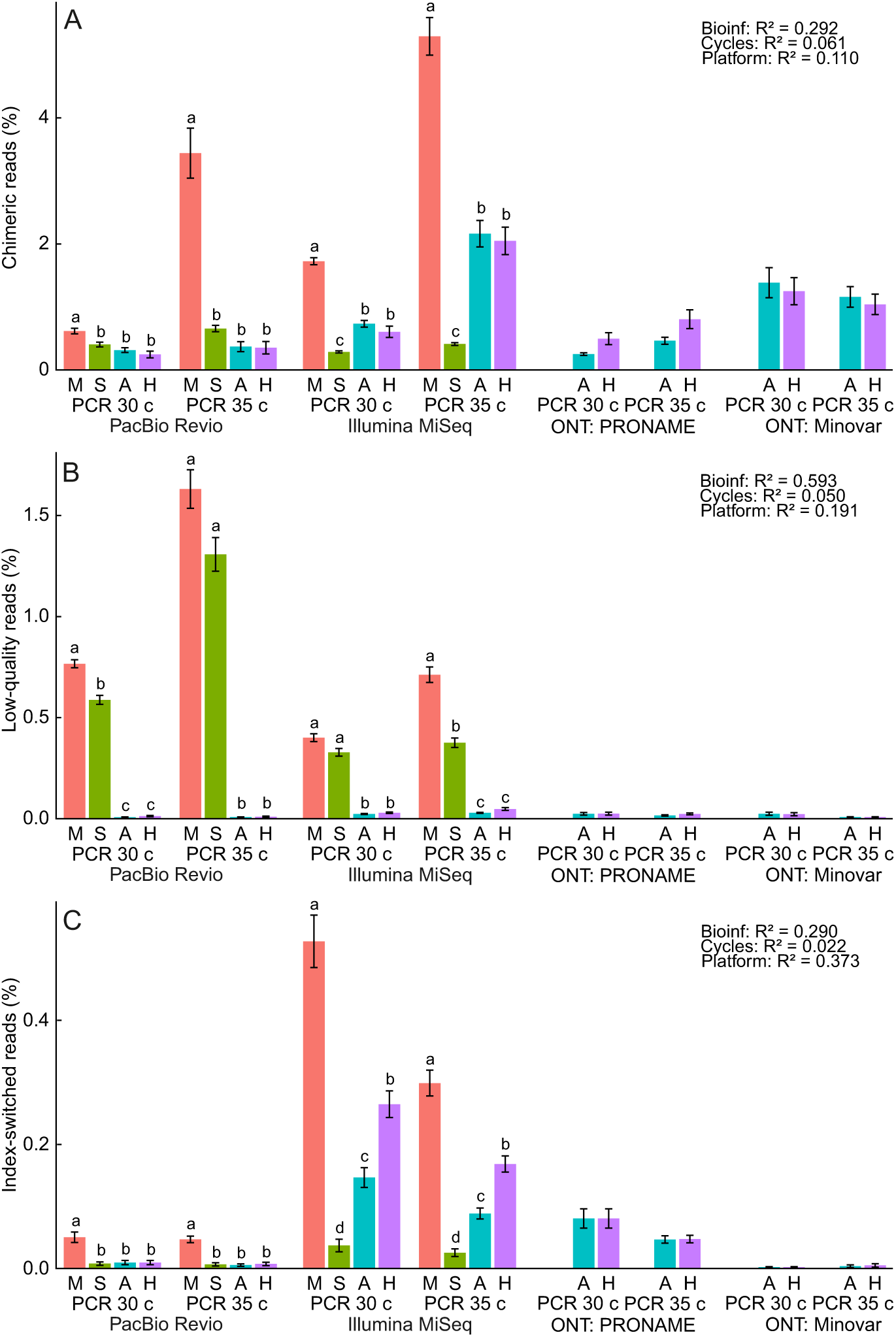
The effects of sequencing platforms, bioinformatics pipelines and PCR cycles on the proportions of A) chimeric reads, B) low-quality reads and C) predicted index-switch artefacts. Bars and errors represent mean±SE. Statistics are based on log-ratio-transformed data. Abbreviations for bioinformatics: M, minimum-filtering; S, standard-filtering; A, ASV; H, hybrid.

The proportion of low-quality sequences was 63.5% higher at PCR35 compared to PCR30 (R^2^=0.001; P<0.001). PacBio yielded a 2.3-fold higher proportion of low-quality reads compared with Illumina (R=0.002; P=0.016), but there were no differences among the ONT-based analysis methods (Figure 5B; Table S4). Minimum-filtered data had the highest proportion of low-quality reads (0.77%), followed by standard-filtering (0.52%), hybrid (0.03%) and ASV (0.02%) approaches (R^2^=0.227; P<0.001).

The proportion of estimated index-switched reads was distinctly highest in the Illumina platform (0.16%), followed by ONT-PRONAME (0.05%), PacBio (0.01%) and ONT-Minovar treatments (<0.01%), with significant differences among all groups (R^2^=0.279; P<0.001), except PacBio and ONT-Minovar (Figure 5C; Table S4). Likewise, index-switched sequences were 11.3-fold, 5.4-fold and 3.0-fold more abundant in the minimum-filtering, hybrid and ASV treatments compared to the standard-filtering treatment (R^2^=0.246; P<0.001).

Differences in the proportions of predicted index-switched OTUs were more pronounced across sequencing platforms. Across all OTUs in Illumina PCR30 soil samples, an average of 53.8%, 26.4%, 14.5% and 3.6% of the minimum-filtered, hybrid, ASV and standard-filtered OTUs, respectively, were estimated to be index-switching artefacts. In comparison, the PCR30 soil samples from PacBio, ONT-Minovar and ONT-PRONAME, respectively, included 0.8%, 0.1% and 7.9% estimated index-switched OTUs.

In the minimum-filtered PCR30 datasets, the number of low-quality, chimeric and index-switched OTUs per sample increased nearly linearly with increasing eukaryote OTU numbers (R range from 0.257 to 0.766). The proportion of low-quality and chimeric OTUs remained similar along the richness gradient, but the proportion of index-switched OTUs declined (PacBio: R=-0.333, P=0.025; Illumina: R=-0.815, P<0.001). In the ASV PCR30 datasets, no change in the proportion of artefacts was evident, and the relationship between chimeric and low-quality OTU numbers with richness occurred only in Illumina and PacBio data, with weaker correlation coefficients (R range from 0.455 to 0.588) than in minimum-filtered data (0.551 to 0.766).

The rarefaction analysis revealed that, while non-artefactual PCR30 OTUs accumulate in a log-linear fashion, up to deepest sequencing, PCR35 OTUs have no or much weaker relationship with sequencing depth (Figure 6A, B). Index-switched OTUs showed a stronger log-linear relationship compared with non-artefactual OTUs (Figure 6G, H). Conversely, both chimeric and low-quality OTUs accumulated linearly with increasing sequencing depth (Figure 6C-F).

**Figure 6.**
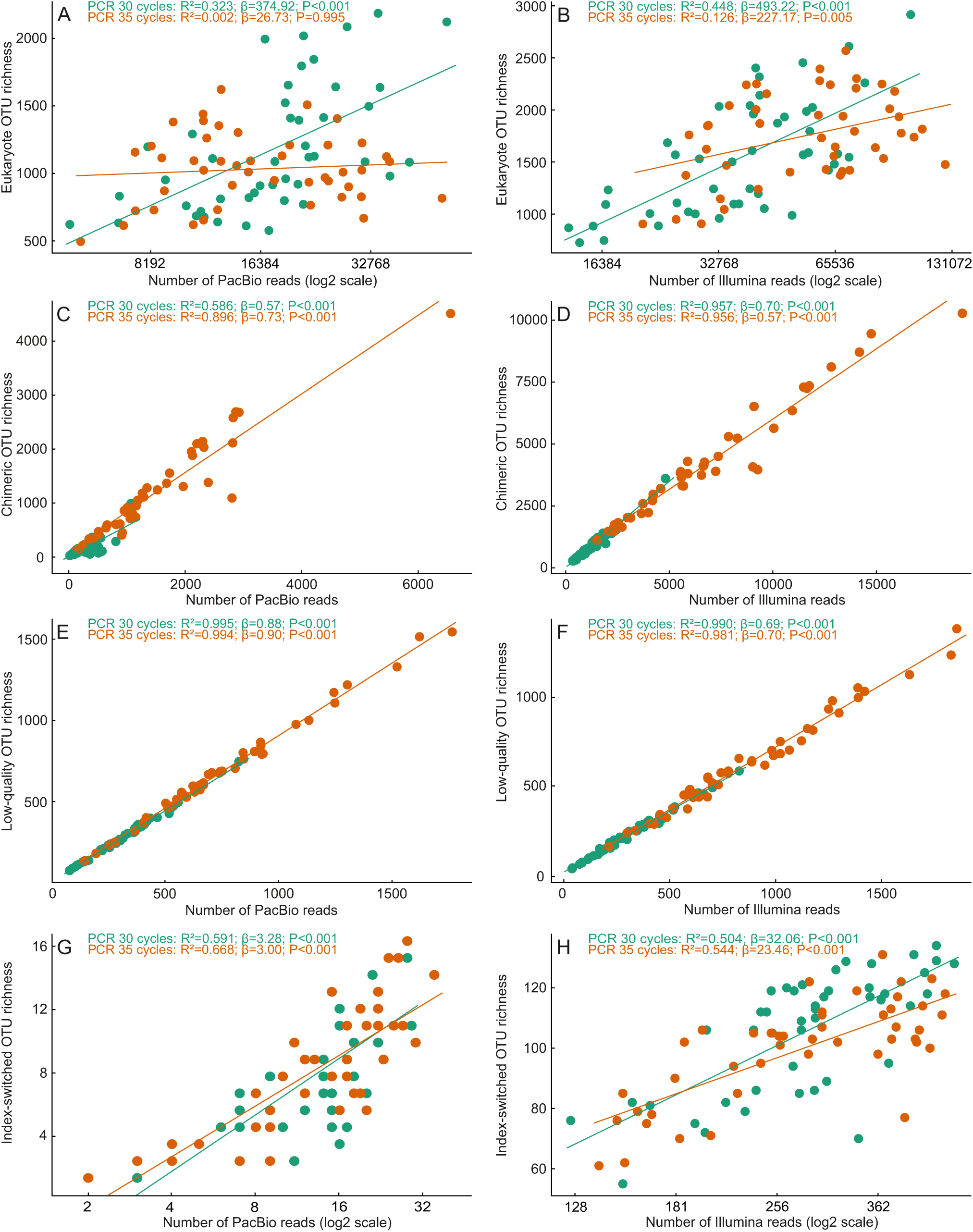
Relative accumulation of A, B) high-quality OTUs; C, D) chimeric OTUs; E, F) low-quality OTUs; and G, H) index-switched OTUs with increasing sequencing depth at PCR30 (green symbols) and PCR35 (orange symbols) in PacBio (left panels) and Illumina (right panels) platforms based on the minimum-filtered dataset. Note differences in the scale of the x-axis.

#### 3.3.3. Sequence length bias

Analysis of soil data revealed biases in OTU composition across sequencing platforms by read length (Figure 7), consistent with the mock community benchmark. The median read length was 776±163, 734±212 and 694±80 bases in PacBio, ONT and Illumina ASV-based datasets, respectively (P>0.001 for all comparisons). While the maximum read length of the amplicon was 2,256 bases for PacBio and 2,686 bases for ONT, the longest recovered Illumina ASV was only 936 bases, partly reflecting the upper limit of 1,000 bases (including primers, tags and the overlap in paired-end assembly) in the 2x500 paired-end mode.

**Figure 7.**
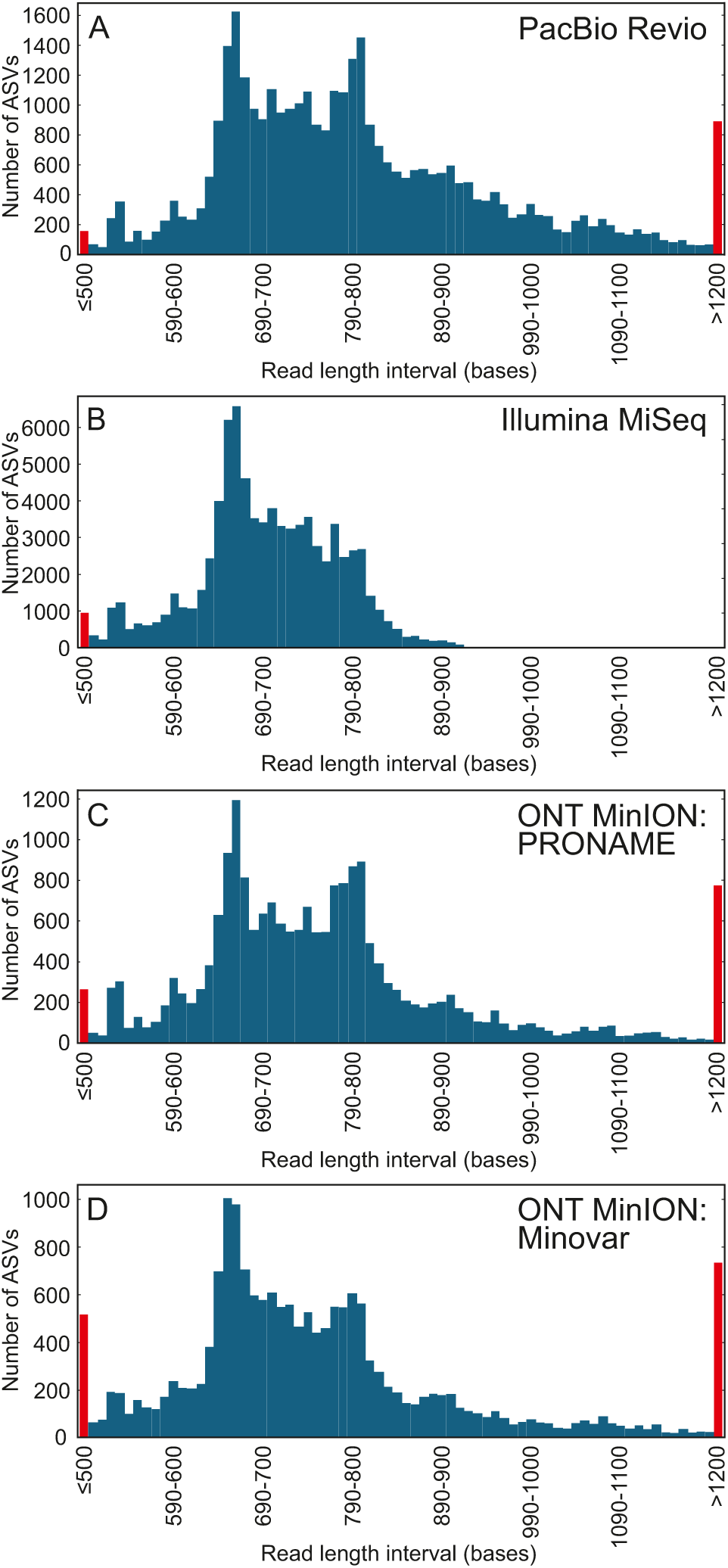
Relative distribution of OTUs by read length across sequencing platforms based on the ASV bioinformatics treatment in A) PacBio Revio, B) Illumina MiSeq 2x500, C) Oxford Nanopore Technologies (ONT) MinION PRONAME and D) ONT MinION Minovar. Red bars indicate total richness below the read length of 500 bases or above 1200 bases. Note the relative loss of Illumina OTUs above amplicon length of ∼750 bases and ONT OTUs above amplicon length of ∼850 bases compared with PacBio, and the relative loss of PacBio OTUs at <500 bases compared with other platforms.

Importantly, 17.4% of PacBio ASVs and 10.7-11.6% ONT ASVs were longer than 936 bases. Among these long PacBio ASVs, around 85% had naturally long ITS1 and/or ITS2 reads, while others were attributable to introns, mainly in the SSU. Taxonomically, these long PacBio ASVs were 4.6-fold and 3.8-fold more common in Metazoa and Rhizaria, respectively, compared with ‘regular’ (<937 bases) ASVs. Conversely, Alveolata, Euglenozoa and Apusomonada were >6-fold less common in the ‘long’ ASV fraction. Fungi contributed to 54.1% of ASVs in the ‘regular’ fraction and 22.6% of ASVs in the ‘long’ fraction. Across all GSMc samples at PCR30, an average of 95.8 ASVs (11.3% of all ASVs) exceeded 936 bases, forming a non-negligible part of the microeukaryote community. In PCR35, 7.4% of ASVs were longer than 936 bases, and the median read length was 15 bases less (P<0.001), indicating an increasing amplicon length bias at surplus cycle numbers.

#### 3.3.4. Comparative analysis of environmental effects

The overall mixed model and most individual models revealed that the more inclusive sampling designs recover greater biodiversity (Table S5). The overall model and most individual models also indicated that forests and grasslands have greater biodiversity than croplands, and there is a significant interaction term between sampling design and land-use type. In particular, forests and especially grasslands had relatively higher biodiversity values than croplands under more inclusive sampling designs (Figure 8). For example, the PacBio standard-filtered PCR30 treatment recovered on average 760±145 (SD) fungal OTUs in croplands, 929±377 OTUs in grasslands and 1144±426 OTUs in forests using the LUCAS design. In croplands, fungal OTU richness increased by 4.1% in the SoilBON design and 6.6% in the GSMc design. In contrast, SoilBON estimates were 39.5% higher for grasslands and 50.5% higher for forests; likewise, GSMc estimates were 119.2% higher for grasslands and by 159.2% higher for forests. Thus, OTU richness estimates differed by 22.2% between grasslands and croplands in the LUCAS design (non-significant), 64.0% in the SoilBON design, and 154.5% in the GSMc design.

**Figure 8.**
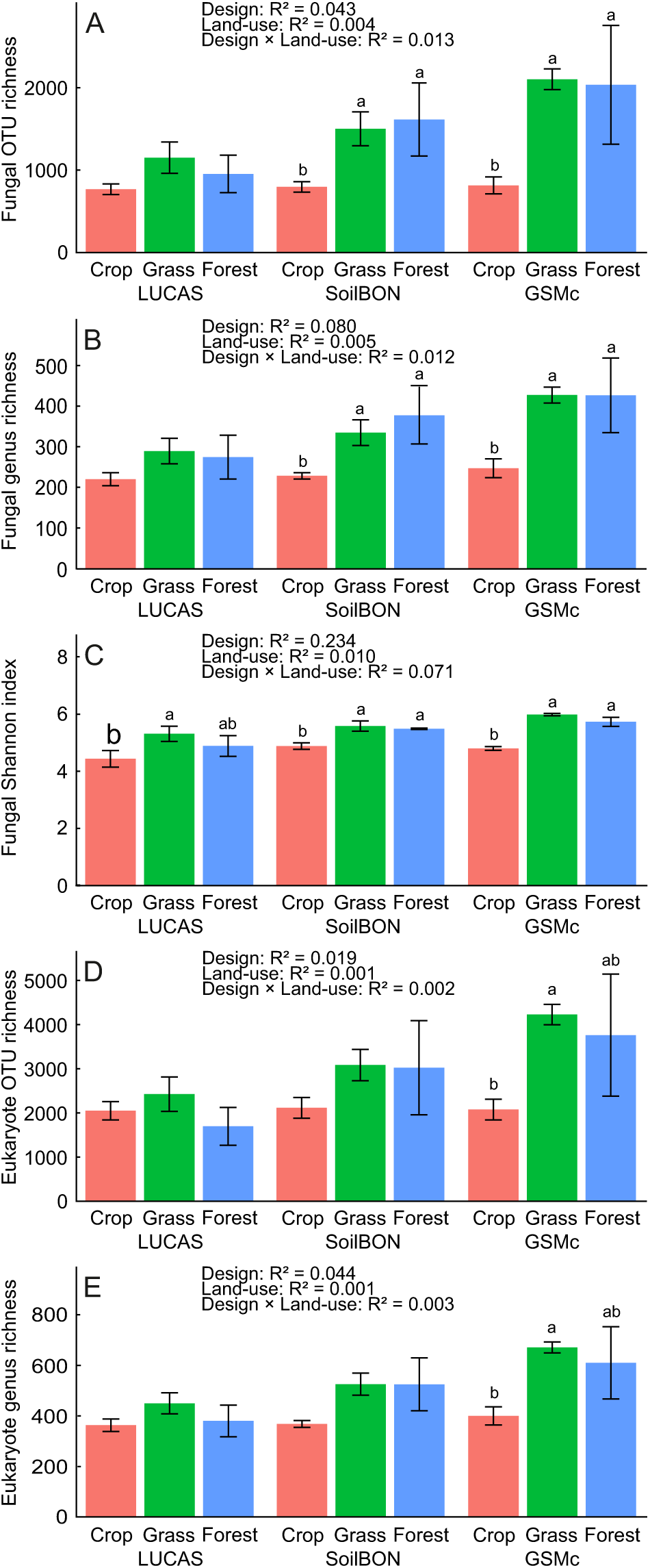
Relative effects of sampling design, land-use and their interaction term in recovering biodiversity values: A) fungal OTU richness, B) fungal genus richness, C) fungal Shannon index, D) eukaryote OTU richness and E) eukaryote genus richness. Data from all sequencing platforms, bioinformatics pipelines and amplicon cycles are pooled for this analysis (n=120 for each group).

The overall mixed model revealed strong effects of sequencing platforms, PCR cycle number and bioinformatics treatment on soil biodiversity variables (Table S5). The Illumina platform recovered the greatest fungal OTU and genus richness, whereas PacBio yielded the highest fungal Shannon index, and ONT-PRONAME found the greatest eukaryote OTU richness. All biodiversity values were significantly greater at PCR30 than PCR35, especially for eukaryote genus richness (R=0.103) and fungal Shannon index (R=0.094). The ASV and hybrid datasets confirmed the overall trends among sequencing platforms (Table S5).

Besides standard biodiversity variables, we compared richness quality, focusing on the number of fungal taxa identical to EUKARYOME reference sequences (Table S5). Across all ASV datasets, PacBio recovered the highest proportion of OTUs identical to sequences in EUKARYOME (average±SD, 61.9+5.0% OTUs), followed by ONT-Minovar (54.8±6.1%), Illumina (47.6±3.4%) and ONT-PRONAME (36.0±4.6% OTUs). In PacBio PCR30 treatment, minimum-filtering (average±SD, 655±281 OTUs) and standard-filtering (613±264) approaches recovered significantly more such identical OTUs than the ASV (265±104) and hybrid (240±92) approaches. Other platforms and the PCR35 data revealed similar trends among bioinformatic treatments.

#### 3.3.5. Methodological comparison for explaining biodiversity

Sequencing platforms, bioinformatics treatments and PCR cycle numbers revealed similar overall trends in explaining biodiversity differences, but differed substantially in their resolution, i.e., effect sizes and statistical significance (Table S6). Across all local models, sampling design, land-use and their interaction respectively explained 19.9±10.6% (SD), 16.2±13.2% and 8.3±3.3% of variation in fungal OTU richness (Figure S1). The sampling design effect was comparable for fungal genus richness (17.8±10.7%), eukaryote OTU richness (19.8±12.6%) and eukaryote genus richness (18.9±13.1%) but lower for fungal Shannon index (12.1±6.4%). The land-use effect was substantially greater for fungal genus richness (33.4±10.1%) and Shannon index (25.8±7.1%) but lower for eukaryote OTU richness (9.0±5.3%) and genus richness (7.8±9.0%). The determination coefficient of the interaction term varied from 0.043±0.021 (fungal Shannon index) to 0.061±0.031 (eukaryote genus richness; Figure S4).

In fungal OTU richness, PCR35 models explained on average 2.2-fold more variance than PCR30 models for the sampling design effect (average R^2^_PCR35_=0.275) and 2.9-fold more variation for the land-use effect (average R^2^ =0.241) but similar magnitude for the interaction (average R^2^ =0.085). Sequencing platforms differed mainly in their estimates of the land-use effect (F=6.91; R^2^=0.147; P=0.003), with Illumina sequencing showing the strongest predictions (average R^2^_Illumina_ =0.265), exceeding those of other platforms by >2-fold.

For fungal genus richness, however, PCR30 provided 1.5-fold stronger estimates for land-use effects than PCR35 and comparable estimates for sampling and interaction effects. For the fungal Shannon index, the PCR30 models explained sampling design 2.3-fold better than the PCR35 models (average R^2^ =0.170).

In eukaryote OTU richness, Illumina models explained >2-fold more variation (average R^2^_Illumina_ =0.347) than models of other platforms for the sampling design effect. For the land-use effect, PacBio (average R^2^=0.112) and Illumina (average R^2^=0.119) explained more than twice the variation as ONT Minovar and PRONAME models. However, the interaction term was best explained by the ONT-PRONAME model, which exceeded models from other platforms by >2-fold. Similar patterns were observed for eukaryote genus richness, except that platforms did not differ in estimates of the interaction effect, and PacBio explained the land-use effect at least 2.9-fold stronger than other platforms (average R^2^_PacBio_=0.162).

The strongest individual models based on the model determination coefficient were Illumina PCR35 standard-filtering and minimum-filtering models for fungal genus richness and eukaryote OTU richness. For fungal OTU richness, Illumina PCR35 hybrid model was equally strong. For eukaryote genus richness, both Illumina and PacBio PCR35 treatments with standard-filtering and minimum-filtering were the strongest. Conversely, for the fungal Shannon index, the ONT-PRONAME PCR30 hybrid and ASV models explained the greatest variation, while ONT-Minovar and ONT-PRONAME PCR35 models performed the weakest. For richness variables, ONT-Minovar and ONT-PRONAME PCR30 models had the lowest determination coefficients.

All sequencing platforms, bioinformatics techniques and PCR cycle numbers explained a comparable amount of fungal community composition (land-use effect: R^2^ range 0.283 to 0.330). The ONT-PRONAME sequencing platform (average R^2^=0.287) and ASV bioinformatics (average R^2^=0.294) tended to yield lower effect sizes than other methods.

## 4. Discussion

### 4.1. Analytical biases

Mock community and soil microbiome data complement each other in revealing multiple critical biases and shortfalls in all sequencing platforms (Hajibabaei et al. 2011; Li et al. 2020; Karst et al. 2021). In our analyses, the mock sample provides near-ground-truth data about chimeric, low-quality and index-switch artefacts in a simplified community, whereas the soil microbiome data offers a replicated, an order of magnitude more diverse real-world situation, where the biases could only be estimated. This combination is important because the mock community identifies the likely technical origin of biases, whereas the soil data show whether these biases are strong enough to affect ecological inference. While estimates of chimeric, low-quality and read-length-related artefacts are consistent, the effect assessments of PCR cycle numbers and index-switches differ. In particular, index-switch rates are greatly underestimated in soil data due to unclear recognition criteria and great differences among sequencing platforms and bioinformatics pipelines. The effects of excess PCR cycles become stronger with increasing biodiversity, where rare taxa may become replaced by chimeras and suppressed by the excess amplification of the abundant taxa.

Illumina 2x500 paired-end sequencing suffers from two main biases: sequence length bias and a high index-switch rate. Although unique symmetric indexes on both primers per sample can minimize index-switching (Schnell et al. 2015; Rodriguez-Martinez et al. 2023), this approach was successful for other platforms but not for Illumina, and the reason remains unclear. Based on mock community samples, we estimate that 3-7% of reads and 15-50% OTUs in the Illumina minimum-filtered and standard-filtered data, and 0.8-2% of sequences and around 15% of taxa in the ASV data, constitute index-switching artefacts. The default UNCROSS thresholds have been developed for ‘regular’ index-switch rates of up to 0.2% of reads.

Therefore, the standard-filtering procedure removes only a small fraction of index-switched OTUs in mock and soil samples, as evidenced by the presence of multiple positive control OTUs in these samples. The ASV and hybrid approaches removed all but the most common index-switched OTUs, suggesting that DADA2 and UNCROSS2, with more stringent abundance thresholds, should be used for analysis of Illumina 2x500 paired-end data, unless technical improvements are implemented in the library preparation or sequencing processes. Regarding the read-length bias, the soil data indicate that amplicons >750 bases are increasingly discriminated against, with >850-bp reads extremely uncommon relative to other platforms, and that an estimated 10-17% of OTUs are lost due to the supplier-determined length limitation of 2x500 paired-end sequencing alone. Based on the mock community with relatively small variation in read length (median±SD: 749±65 bases), we calculate that amplicon length bias accounts for 15-26% of variation in the relative abundance of OTUs.

This bias is expected to be much stronger in the more heterogeneous soil data (median sequence length 776±163 bases), with additional losses of taxa that exhibit ITS sequences >750-850 bases. Our analysis shows that the taxa with naturally long ITS amplicons, including many metazoans and cercozoans, and those with SSU 3’ proximal introns (commonly ascomycetes and nematodes), become enormously downsampled. Conversely, taxa with predominantly short ITS sequences, such as apicomplexans, euglenozoans and apusomonads are favoured, reinforcing the read-length bias (Kanagawa 2003). However, selecting PCR primers closer to the ITS region should partially ameliorate this bias because of the resulting 10-25% shorter amplicons.

PacBio serves as a benchmark platform for long-read metabarcoding analyses due to the high quality of HiFi reads produced by CCS (Jamy et al. 2020; Tedersoo et al. 2021a; Overgaard et al. 2024). PacBio produced the highest proportion of OTUs that perfectly matched sequences in the EUKARYOME database, confirming the highest data quality of HiFi reads. However, our analysis also indicates that HiFi data can include large numbers of reads with substandard quality masked by high Phred scores. In our soil data, the PacBio minimum-filtered and standard-filtered treatments produced the highest numbers of low-quality reads and OTUs per sample. With multiple indels, these reads commonly formed low-abundance satellite OTUs in the periphery of common OTUs (Frøslev et al. 2017; Bonin et al. 2023). Furthermore, PacBio revealed around 30% lower richness of apicomplexans and ciliates, both with relatively short ITS regions, compared to Illumina and ONT platforms. Discrimination against >500-base reads accords with known limitations of PacBio Revio chemistry (Pacific Biosciences 2025).

Due to the low initial read quality, the main analytical shortfall of ONT is the loss of a large proportion of reads during demultiplexing and stringent quality-filtering as implemented in PRONAME. The ONT Minovar data slightly but significantly exceeded the ASV quality of Illumina sequencing data, indicating that recent versions of read polishing software perform well. The high proportion of low-quality reads observed in the mock community ASV data was nearly eliminated by the hybrid approach. Minovar requires at least 5 reads per high-similarity cluster within each sample to build high-quality consensus sequences; thus, all rare OTUs with <5 reads per sample are automatically eliminated, and a large proportion of OTUs with <10 reads are lost due to failed consensus building if these reads belong to different species. Conversely, PRONAME includes an initial quality-score-based filtering step that removes a larger proportion of reads, leaving less data for consensus building. The need to retain more clusters, supported by a low number of reads, very likely contributed to the greater proportion of index-switched reads observed with PRONAME, and underscores the potential value of removing very rare entities from ONT data. At present, we recommend Minovar analysis of ONT ITS amplicon data using the hybrid approach. The analysis would benefit further from removing extra-low-quality reads.

All three bioinformatics pipelines greatly improved the data quality of all sequencing platform outputs, which initially contained low-quality and chimeric reads and index-switch artefacts. Standard-filtering, involving additional chimera and low-quality data removal and index-switch control, efficiently removes all types of artefacts, except index-switch artefacts in Illumina sequencing. The ASV and hybrid methods are similarly efficient at removing all types of artefacts in PacBio and ONT-Minovar data. Still, they are less efficient at removing chimeras and index-switches in Illumina and ONT-PRONAME data. While standard filtering removes only 0.7-1.1% of genera, mostly due to index switches, ASV and hybrid methods lose 42-49% of genera and 50-75% of expected high-quality OTUs per sample. This argues against the common misconception that ASV-based datasets are closer to saturated sequencing depth, because many low-abundant taxa have been removed (Joos et al. 2020).

Excess PCR cycles reduced concordance among sequencing platforms, primarily by accentuating amplicon-length bias and introducing chimeric and low-quality reads and OTUs. Specifically, the PCR35 treatment yielded on average 2.7-fold more chimeric and 2.2-fold more low-quality reads than the PCR30 treatment in the soil data. Multiple studies have shown that excess numbers of PCR cycles produce more chimeras (D’Amore et al. 2016; Gohl et al. 2016; Wang & Wang 1996) and other artefacts (Castaño et al. 2020). Consistent with previous research (Ho et al. 2021), higher cycle numbers also reduced fungal Shannon diversity in soil communities, thereby weakening the semiquantitative properties of HTS and suggesting that multiple rare OTUs may be outcompeted during excess amplification and remain undetected.

For practical full-length ITS metabarcoding, the preferred methods depend on the analytical goal. PacBio with standard OTU-based filtering is the most balanced option for biodiversity surveys, where retention of rare taxa and broad amplicon-length coverage are important. If the primary aim is deep sequencing of a constant-length marker, Illumina 2x500 sequencing with ASV-based or otherwise stringent filtering could be used. The ONT platform is the most useful when rapid, in-house long-read sequencing is required, or dominant community members or certain indicator taxa are of primary interest; here, Minovar-based hybrid clustering is preferable over ASVs, because rare low-abundance variants and low-quality consensus sequences remain problematic.

### 4.2. Ecological insights

The comparative analysis of soil samples indicates that sequencing platforms, PCR cycle numbers and bioinformatics treatments differ in the relative recovery of soil biodiversity variables. After accounting for sequencing depth, Illumina yielded the greatest fungal richness, whereas PacBio recovered the highest fungal Shannon index. Standard-filtering outperformed ASV and hybrid approaches in recovering higher OTU and genus-level richness by retaining rare but high-quality OTUs (see above).

The technical options differed greatly in their ability to recover the effects of sampling design, land use and their interaction term on various fungal and eukaryotic biodiversity variables. Contrary to our predictions, models built on expectedly noisier data yielded stronger effects of experimental variables on biodiversity than models built on better-filtered datasets, often explaining 2-fold more variation. These findings suggest that greater accumulation of errors at higher levels of biodiversity may emphasize, rather than blur, the ecological differences among experimental factors. Furthermore, chimeric and low-quality OTUs accumulate linearly with increasing sequencing depth, whereas high-quality OTUs and index-switches accumulate logarithmically (Figure 6). The relatively poor performance of ONT-based methods, as well as ASV and hybrid pipelines in general, indicates that these methods lose too many rare biological OTUs that underpin overall biodiversity and multifunctionality estimates (Zhang et al. 2022).

Supporting our ecological hypothesis, most models indicated that more representative sampling (i.e., a larger pool of subsamples) enhanced the detection of differences in soil biodiversity among land-use types. These findings suggest that spatially limited sampling designs, such as LUCAS, are more suitable for croplands than for natural grasslands and forests, due to both the spatial scale of sampling and spatial heterogeneity of communities. Given the long-term, multi-seasonal disturbance, croplands have become highly homogeneous at the field scale and across larger geographic scales (Labouyrie et al. 2023; Peng et al. 2024). Because of the disturbances and high fertility, croplands harbour copiotrophic microbiota characterised by efficient spore dispersal, rapid growth and reproductive cycles (He et al. 2025). Grasslands and, particularly, forests develop microsites that differ in physicochemical conditions and support perennial plants that create locally specific rhizosphere environments. Such fine-scale environmental variability promotes biodiversity in many organism groups via niche differentiation (Lindahl et al. 2007; Peršoh et al. 2018). When projecting to the pan-European LUCAS soil survey, our regional-scale study cannot fully predict potential sampling design effects in other bioclimatic zones.

### 4.3. Limitations

The main limitation is the lack of replication of the sequencing process, although this is common to a vast majority of such comparative studies (but see Sun et al. 2020). We also did not compare the broader array of Illumina library preparation methods that may affect artefact formation (Bronner & Quail 2019; Sato et al. 2019). Comparisons among sequencing platforms are complicated by the need for platform-specific preprocessing, especially for ONT data, where consensus generation and polishing are indispensable. A fully identical preprocessing workflow would therefore be technically unrealistic and would not represent best-practice use of each platform, but this also means that some observed differences reflect platform-pipeline combinations rather than sequencing platforms alone. Our mock community consists exclusively of fungal taxa, all with near-average ITS sequence lengths, and no information is available on the number of ITS copies per specimen or DNA integrity. These features do not allow certain quantitative estimates of biases, and caution should be taken when extrapolating to other taxa and markers.

## 5. Conclusion

Our comparative analyses show that for full-length ITS metabarcoding, PacBio is preferred over Illumina and ONT because of relatively low artefact rates (excluding low-quality reads) and better demultiplexing equality, with comparable sequencing costs per read. For PacBio data, the relatively commonly used OTU-based quality-filtering approaches outperform the DADA2 ASV and hybrid approaches, which lose too much of the rare biosphere and, hence, underestimate biodiversity. Given the high error rates in ONT and index-switch issues in Illumina MiSeq 2x500 mode, these data should be processed with stringent quality-filtering options that eliminate rare OTUs in each sample. We also demonstrate that Minovar is an efficient software for ONT data quality-filtering that approximates common sequence variants at the SNP level in mock community and eDNA data, with optimal performance using the hybrid approach. The same technical choices evaluated in the mock community affect biodiversity estimates in soil communities, but they do not overturn the central ecological conclusion: more representative sampling designs improve the detection of land-use effects on soil biodiversity.

## Ethics statement

Our work follows all ethical and research integrity standards.

## Acknowledgements

We thank three anonymous reviewer for their constructive criticism. We are indebted to R. Puusepp for performing molecular analyses. We acknowledge the Norwegian Sequencing Centre (Oslo) and Illumina Solutions Centre (Berlin) for the sequencing services. This study received funding from the ERC-Advanced grant 101200758 (PhylFun) and Centre of Excellence AgroCropFuture (TK200) and Research Council of Finland (Decision number 362828). M.C. thanks the China Scholarship Council for a doctorate grant at Tartu University.

## Data Availability and Benefit sharing

Raw sequencing reads generated on all three sequencing platforms have been deposited in the European Nucleotide Archive (ENA) under the project accession PRJEB108994 (sample accession numbers ERS29406352 - ERS29406397; sequence accession numbers ERR16773363 - ERR16773639). Sequences and metadata of mock community specimens are available in UNITE (Table S1).

The Source code is available at https://github.com/Mycology-Microbiology-Center/fullITS-multiplatform-eval. The Minovar pipeline v1.0 is archived at Zenodo at https://zenodo.org/records/18803397.

Benefits from this research accrue from the sharing of our data and results on public databases as described above.

## Author Contributions

L.T. designed research, analysed data and wrote the paper, with contributions from all co-authors; M.P. generated and used the Minovar software; M.C. performed GLM and multivariate modelling; B.D. analysed ONT data using PRONAME; V.M. and S.A. performed bioinformatics analyses. I.S. generated Sanger sequencing data for the mock community.

## Supplementary materials

**Figure S1.** Relationship of OTU abundance and its ITS amplicon length based on the ASV data across sequencing platform and PCR cycle number combinations in the mock community.

**Figure S2.** Pearson correlations of PacBio-based fungal (top rows) and eukaryote (bottom rows) richness residuals with the corresponding richness residuals from Illumina MiSeq (left panels), Oxford Nanopore Minion (ONT) PRONAME (central panels) and ONT Minovar (right panels).

**Figure S3.** Pearson correlations of fungal (top rows) and eukaryote (bottom rows) OTU richness residuals based on 30 and 35 PCR cycles.

**Figure S4.** Relative effect sizes of sequencing platforms (left panels), bioinformatics pipelines (central panels) and amplicon cycle numbers (right panels) for explaining sampling design, land-use and their interaction effects on A) fungal OTU richness, B) fungal genus richness, C) fungal Shannon index, D) eukaryote OTU richness and E) eukaryote genus richness.

**Table S1.** Relative efficiency of sequencing platforms and bioinformatics pipelines in recovering species in the mock community.

**Table S2.** Details of soil samples used in the analysis.

**Table S3.** Rank distribution of 50 most abundant taxa derived from soil samples across sequencing platforms and bioinformatics treatments.

**Table S4.** Mixed models assessing the effects of sequencing platforms, bioinformatics pipelines and PCR cycle numbers on recovering artefactual reads.

**Table S5.** Mixed models assessing the effects of sequencing platforms, bioinformatics pipelines and PCR cycle numbers on recovering fungal and eukaryote biodiversity.

**Table S6.** Relative performance of sequencing platforms, bioinformatics pipelines and PCR cycle numbers in recovering the effects of sampling design, land use and their interaction on fungal and eukaryote biodiversity variables.

## References

Anslan, S., and L. Tedersoo. 2015. “Performance of cytochrome c oxidase subunit I (COI), ribosomal DNA Large Subunit (LSU) and Internal Transcribed Spacer 2 (ITS2) in DNA barcoding of Collembola.” European Journal of Soil Biology 69: 1–7.

Antil, S., J.S. Abraham, S. Sripoorna, et al. 2023. “DNA barcoding, an effective tool for species identification: A review.” Molecular Biology Reports 50: 761–775.

Arslan, S., F.J. Garcia, M. Guo, et al. 2024. “Sequencing by avidity enables high accuracy with low reagent consumption.” Nature Biotechnology 42: 132–138.

Aslani, F., M. Bahram, S. Geisen, et al. 2024. “Land use intensification homogenizes soil protist communities and alters their diversity across Europe.” Soil Biology and Biochemistry 195: 109459.

Bates, D., M. Mächler, B. Bolker, and S. Walker. 2015. “Fitting Linear Mixed-Effects Models Using lme4." Journal of Statistical Software 67: 1–48.

BengtssonLJPalme, J., M. Ryberg, M. Hartmann, et al. 2013. “Improved software detection and extraction of ITS1 and ITS2 from ribosomal ITS sequences of fungi and other eukaryotes for analysis of environmental sequencing data.” Methods in Ecology and Evolution 4: 914–919.

Bonin, A., A. Guerrieri, and G.F. Ficetola. 2023. “Optimal sequence similarity thresholds for clustering of molecular operational taxonomic units in DNA metabarcoding studies.” Molecular Ecology Resources 23: 368–381.

Bronner, I.F., and M.A. Quail. 2019. “Best Practices for Illumina Library Preparation.” Current Protocols in Human Genetics 102: e86.

Callahan, B.J., P.J. McMurdie, M.J. Rosen, et al. 2016. “DADA2: High-resolution sample inference from Illumina amplicon data.” Nature Methods 13: 581–583.

Camacho, C., G. Coulouris, V. Avagyan, et al. 2009. “BLAST+: Architecture and applications.” BMC Bioinformatics 10: 421.

Carlsen, T., A.B. Aas, D. Lindner, et al. 2012. “Don’t make a mista(g)ke: Is tag switching an overlooked source of error in amplicon pyrosequencing studies?” Fungal Ecology 5: 747–749.

Castaño, C., A. Berlin, M. Brandström Durling, et al. 2020. “Optimized metabarcoding with Pacific biosciences enables semiLJquantitative analysis of fungal communities.” New Phytologist 228: 1149–1158.

Chen, M., O. Dulya, V. Mikryukov, et al. 2026. “Sampling Design and Sample Processing Affect Soil Biodiversity Assessments.” Molecular Ecology Resources 26: e70113.

D’Amore, R., U.Z. Ijaz, M. Schirmer, et al. 2016. “A comprehensive benchmarking study of protocols and sequencing platforms for 16S rRNA community profiling.” BMC Genomics 17: 55.

Deiner, K., H.M. Bik, E. Mächler, et al. 2017. “Environmental DNA metabarcoding: Transforming how we survey animal and plant communities.” Molecular Ecology 26: 5872–5895.

Di Tommaso, P., M. Chatzou, E.W. Floden, et al. 2017. “Nextflow enables reproducible computational workflows.” Nature Biotechnology 35: 316–319.

Dubois, B., M. Delitte, S. Lengrand, et al. 2024. “PRONAME: A user-friendly pipeline to process long-read nanopore metabarcoding data by generating high-quality consensus sequences.” Frontiers in Bioinformatics 4: 1483255.

Edgar, R.C. 2010. “Search and clustering orders of magnitude faster than BLAST.” Bioinformatics 26: 2460–2461.

Edgar, R.C. 2016. “UNOISE2: Improved error-correction for Illumina 16S and ITS amplicon sequencing.” bioRxiv 2016: 081257.

Edgar, R.C. 2018. “UNCROSS2: Identification of cross-talk in 16S rRNA OTU tables.” bioRxiv 2018: 400762.

Edgar, R.C., B.J. Haas, J.C. Clemente, et al. 2011. “UCHIME improves sensitivity and speed of chimera detection.” Bioinformatics 27: 2194–2200.

Edwards, D.P., and G.R. Cerullo. 2024. “Biodiversity is central for restoration.” Current Biology 34: R371–R379.

Froger, C., E. Tondini, D. Arrouays, et al. 2024. “Comparing LUCAS Soil and national systems: Towards a harmonized European Soil monitoring network.” Geoderma 449: 117027.

Frøslev, T.G., R. Kjøller, H.H. Bruun, et al. 2017. “Algorithm for post-clustering curation of DNA amplicon data yields reliable biodiversity estimates.” Nature Communications 8: 1188.

Gao, Y., Y. Liu, Y. Ma, et al. 2021. “abPOA: An SIMD-based C library for fast partial order alignment using adaptive band.” Bioinformatics 37: 2209–2211.

Garrison, E., and G. Marth. 2012. “Haplotype-based variant detection from short-read sequencing (Version 2).” arXiv 1207: 3907.

Gohl, D.M., P. Vangay, J. Garbe, et al. 2016. “Systematic improvement of amplicon marker gene methods for increased accuracy in microbiome studies.” Nature Biotechnology 34: 942–949.

Guerra, C.A., R.D. Bardgett, L. Caon, et al. 2021. “Tracking, targeting, and conserving soil biodiversity.” Science 371: 239–241.

Hajibabaei, M., S. Shokralla, X. Zhou, et al. 2011. “Environmental Barcoding: A Next-Generation Sequencing Approach for Biomonitoring Applications Using River Benthos.” PLoS ONE 6: e17497.

He, D., Z. Dai, S. Cheng, et al. 2025. “Microbial life-history strategies and genomic traits between pristine and cropland soils.” mSystems 10: e00178–25.

Ho, M., D. Moon, M. Pires-Alves, et al. 2021. “Recovery of microbial community profile information hidden in chimeric sequence reads.” Computational and Structural Biotechnology Journal 19: 5126–5139.

Illumina. 2025. “The New Standard in Benchtop Sequencing Speed and Simplicity.” Available at: https://emea.illumina.com/systems/sequencing-platforms/miseq-i100.html

Jamy, M., R. Foster, P. Barbera, et al. 2020. “LongLJread metabarcoding of the eukaryotic rDNA operon to phylogenetically and taxonomically resolve environmental diversity.” Molecular Ecology Resources 20: 429–443.

Joos, L., S. Beirinckx, A. Haegeman, et al. 2020. “Daring to be differential: Metabarcoding analysis of soil and plant-related microbial communities using amplicon sequence variants and operational taxonomical units.” BMC Genomics 21: 733.

Kanagawa, T. 2003. “Bias and artifacts in multitemplate polymerase chain reactions (PCR).” Journal of Bioscience and Bioengineering 96: 317–323.

Karst, S.M., R.M. Ziels, R.H. Kirkegaard, et al. 2021. “High-accuracy long-read amplicon sequences using unique molecular identifiers with Nanopore or PacBio sequencing.” Nature Methods 18: 165–169.

Kauserud, H. 2023. “ITS alchemy: On the use of ITS as a DNA marker in fungal ecology.” Fungal Ecology 65: 101274.

Köninger, J., C. Ballabio, P. Panagos, et al. 2023. “Ecosystem type drives soil eukaryotic diversity and composition in Europe.” Global Change Biology 29: 5706–5719.

Kumar, K.R., M.J. Cowley, and R.L. Davis. 2019. “Next-Generation Sequencing and Emerging Technologies.” Seminars in Thrombosis and Hemostasis 45: 661–673.

Labouyrie, M., C. Ballabio, F. Romero, et al. 2023. “Patterns in soil microbial diversity across Europe.” Nature Communications 14: 3311.

Li, H. 2021. “New strategies to improve minimap2 alignment accuracy.” Bioinformatics 37: 4572–4574.

Li, S., Y. Deng, Z. Wang, et al. 2020. “Exploring the accuracy of ampliconLJbased internal transcribed spacer markers for a fungal community.” Molecular Ecology Resources 20: 170–184.

Lindahl, B.D., K. Ihrmark, J. Boberg, et al. 2007. “Spatial separation of litter decomposition and mycorrhizal nitrogen uptake in a boreal forest.” New Phytologist 173: 611–620.

Loit, K., K. Adamson, M. Bahram, et al. 2019. “Relative Performance of MinION (Oxford Nanopore Technologies) versus Sequel (Pacific Biosciences) Third-Generation Sequencing Instruments in Identification of Agricultural and Forest Fungal Pathogens.” Applied and Environmental Microbiology 85: e01368–19.

Prous, M., and V. Mikryukov. 2026. “Minovar (Version 1.0).” Zenodo. 10.5281/zenodo.18803397

Martin, M. 2011. “Cutadapt removes adapter sequences from high-throughput sequencing reads.” EMBnet Journal 17: 10.

Mikryukov, V. 2025. “phredsort: A command-line tool for sorting sequences in a FASTQ file by their quality scores (Version 1.4.0).” Zenodo. 10.5281/zenodo.14395125

Mikryukov, V., S. Anslan, and L. Tedersoo. 2026. “NextITS: A pipeline for metabarcoding eukaryotes with full-length ITS sequenced with PacBio (Version 1.1.0).” Zenodo. 10.5281/zenodo.15074881

Orgiazzi, A., C. Ballabio, P. Panagos, et al. 2018. “LUCAS Soil, the largest expandable soil dataset for Europe: A review.” European Journal of Soil Science 69: 140–153.

Overgaard, C.K., M. Jamy, S. Radutoiu, et al. 2024. “Benchmarking longLJread sequencing strategies for obtaining ASVLJresolved rRNA operons from environmental microeukaryotes.” Molecular Ecology Resources 24: e13991.

Pacific Biosciences. 2025. “Full-Length HiFi Sequencing of Amplicons.” Available at: https://www.pacb.com/wp-content/uploads/Application-Brief-Targeted-sequencing-for-amplicons-best-practices.pdf

Pawlowski, J., S. Audic, S. Adl, et al. 2012. “CBOL Protist Working Group: Barcoding Eukaryotic Richness beyond the Animal, Plant, and Fungal Kingdoms.” PLoS Biology 10: e1001419.

Peng, Z., X. Qian, Y. Liu, et al. 2024. “Land conversion to agriculture induces taxonomic homogenization of soil microbial communities globally.” Nature Communications 15: 3624.

Peršoh, D., N. Stolle, A. Brachmann, et al. 2018. “Fungal guilds are evenly distributed along a vertical spruce forest soil profile while individual fungi show pronounced niche partitioning.” Mycological Progress 17: 925–939.

R Core Team. 2023. “R: A Language and Environment for Statistical Computing.” R Foundation for Statistical Computing. Available at: https://www.r-project.org/

RodriguezLJMartinez, S., J. Klaminder, M.A. Morlock, et al. 2023. “The topological nature of tag jumping in environmental DNA metabarcoding studies.” Molecular Ecology Resources 23: 621–631.

Rognes, T., T. Flouri, B. Nichols, et al. 2016. “VSEARCH: A versatile open source tool for metagenomics.” PeerJ 4: e2584.

Santoferrara, L., F. Burki, S. Filker, et al. 2020. “Perspectives from Ten Years of Protist Studies by HighLJThroughput Metabarcoding.” Journal of Eukaryotic Microbiology 67: 612–622.

Sato, M.P., Y. Ogura, K. Nakamura, et al. 2019. “Comparison of the sequencing bias of currently available library preparation kits for Illumina sequencing of bacterial genomes and metagenomes.” DNA Research 26: 391–398.

Schnell, I. B., K. Bohmann, and M.T.P. Gilbert. 2015. “Tag jumps illuminated – reducing sequenceLJtoLJsample misidentifications in metabarcoding studies.” Molecular Ecology Resources 15: 1289–1303.

Schoch, C.L., K.A. Seifert, S. Huhndorf, et al. 2012. “Nuclear ribosomal internal transcribed spacer (ITS) region as a universal DNA barcode marker for Fungi.” Proceedings of the National Academy of Sciences 109: 6241–6246.

Shaw, J., C. Boucher, Y. Yu, et al. 2025. “Long-read reconstruction of many diverse haplotypes with devider.” Genome Research 35: 2637–2649.

Shen, W., B. Sipos, and L. Zhao. 2024. “SeqKit2: A Swiss army knife for sequence and alignment processing.” iMeta 3: e191.

Sievers, F., A. Wilm, D. Dineen, et al. 2011. “Fast, scalable generation of highLJquality protein multiple sequence alignments using Clustal Omega.” Molecular Systems Biology 7: 539.

Smith, M.E., G.W. Douhan, and D.M. Rizzo. 2007. “Intra-specific and intra-sporocarp ITS variation of ectomycorrhizal fungi as assessed by rDNA sequencing of sporocarps and pooled ectomycorrhizal roots from a Quercus woodland.” Mycorrhiza 18: 15–22.

Stoler, N., and A. Nekrutenko. 2021. “Sequencing error profiles of Illumina sequencing instruments.” NAR Genomics and Bioinformatics 3: lqab019.

Sun, X., J. Wang, C. Fang, et al. 2020. “Efficient and stable metabarcoding sequencing from DNBSEQ-G400 sequencer examined by large fungal community analysis.” bioRxiv 2020: 2020.07.02.185710

Taberlet, P., E. Coissac, F. Pompanon, et al. 2012. “Towards nextLJgeneration biodiversity assessment using DNA metabarcoding.” Molecular Ecology 21: 2045–2050.

Tedersoo, L., M. Albertsen, S. Anslan, et al. 2021. “Perspectives and Benefits of High-Throughput Long-Read Sequencing in Microbial Ecology.” Applied and Environmental Microbiology 87: e00626–21.

Tedersoo, L., M. Bahram, L. Zinger, et al. 2022. “Best practices in metabarcoding of fungi: From experimental design to results.” Molecular Ecology 31: 2769–2795.

Tedersoo, L., M.S. Hosseyni Moghaddam, V. Mikryukov, et al. 2024. “EUKARYOME: The rRNA gene reference database for identification of all eukaryotes.” Database 2024: baae043.

Tedersoo, L., V. Mikryukov, S. Anslan, et al. 2021. “The Global Soil Mycobiome consortium dataset for boosting fungal diversity research.” Fungal Diversity 111: 573–588.

Tilman, D., F. Isbell, and J.M. Cowles. 2014. “Biodiversity and Ecosystem Functioning.” Annual Review of Ecology, Evolution, and Systematics 45: 471–493.

Varusk, S., K. Sammet, M. Ariyan, et al. 2025. “DNA metabarcoding of mites from small soil samples: Limited agreement with morphological identifications but improved results from long-read sequencing.” PeerJ 13: e20205.

Wang, G.C.Y., and Y. Wang. 1996. “The frequency of chimeric molecules as a consequence of PCR co-amplification of 16S rRNA genes from different bacterial species.” Microbiology 142: 1107–1114.

Wick, R.R. 2023. “ONT accuracy vs depth update (Version v1.0.0).” Zenodo. 10.5281/zenodo.10073815

Zhang, Z., Y. Lu, G. Wei, et al. 2022. “Rare Species-Driven Diversity–Ecosystem Multifunctionality Relationships are Promoted by Stochastic Community Assembly.” mBio 13: e00449–22.

Zhang, T., M. Jiang, H. Li, et al. 2025. “Computational Tools and Resources for Long-read Metagenomic Sequencing Using Nanopore and PacBio.” Genomics, Proteomics & Bioinformatics 23: qzaf075.

