## Supplementary material for "Benchmarking full-length ITS metabarcoding across Illumina 2x500, PacBio, and Oxford Nanopore sequencing using mock and soil communities": Figures S1-S4.pdf

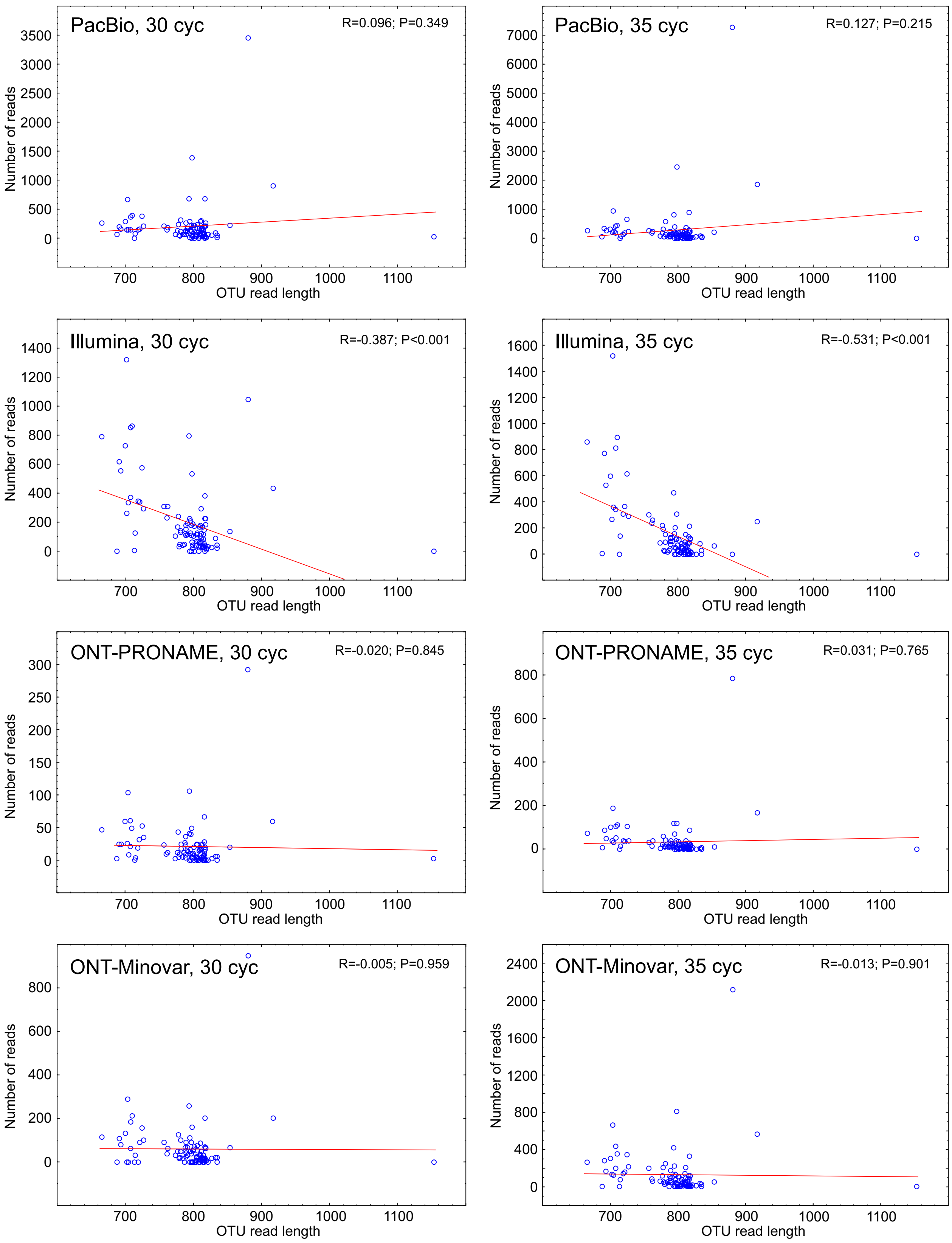


**Figure S1.** Relationship of OTU abundance and its ITS amplicon length based on the ASV data across sequencing platform and PCR cycle number combinations in the mock community. The solid line depicts linear regression line. Pearson R and corresponding P-values are indicated.

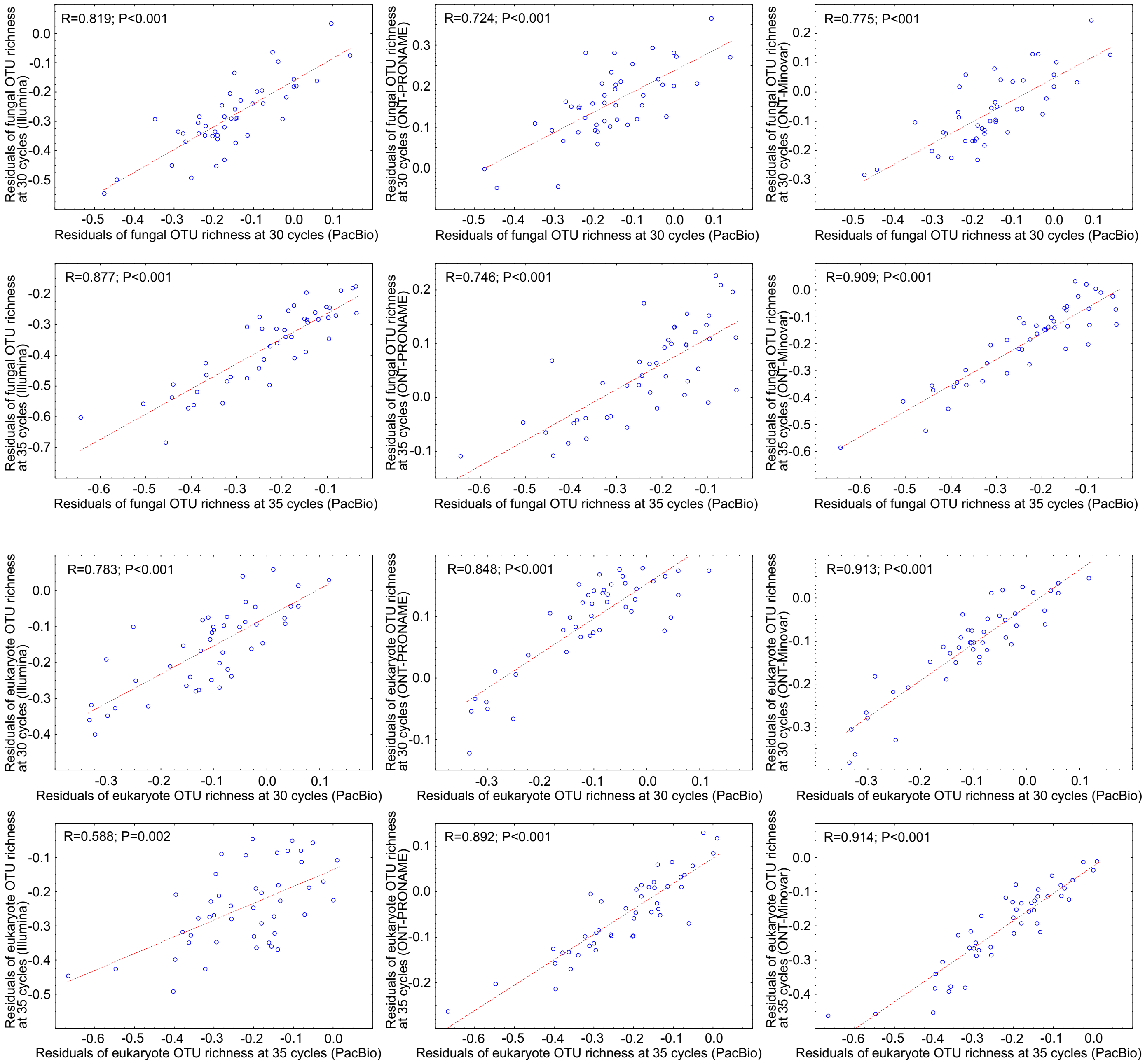


**Figure S2**. Pearson correlations of PacBio-based fungal (top rows) and eukaryote (bottom rows) richness residuals with the corresponding richness residuals from Illumina MiSeq (left panels), Oxford Nanopore Minion (ONT) PRONAME (central panels) and ONT Minovar (right panels). Dashed lines indicate linear fit.

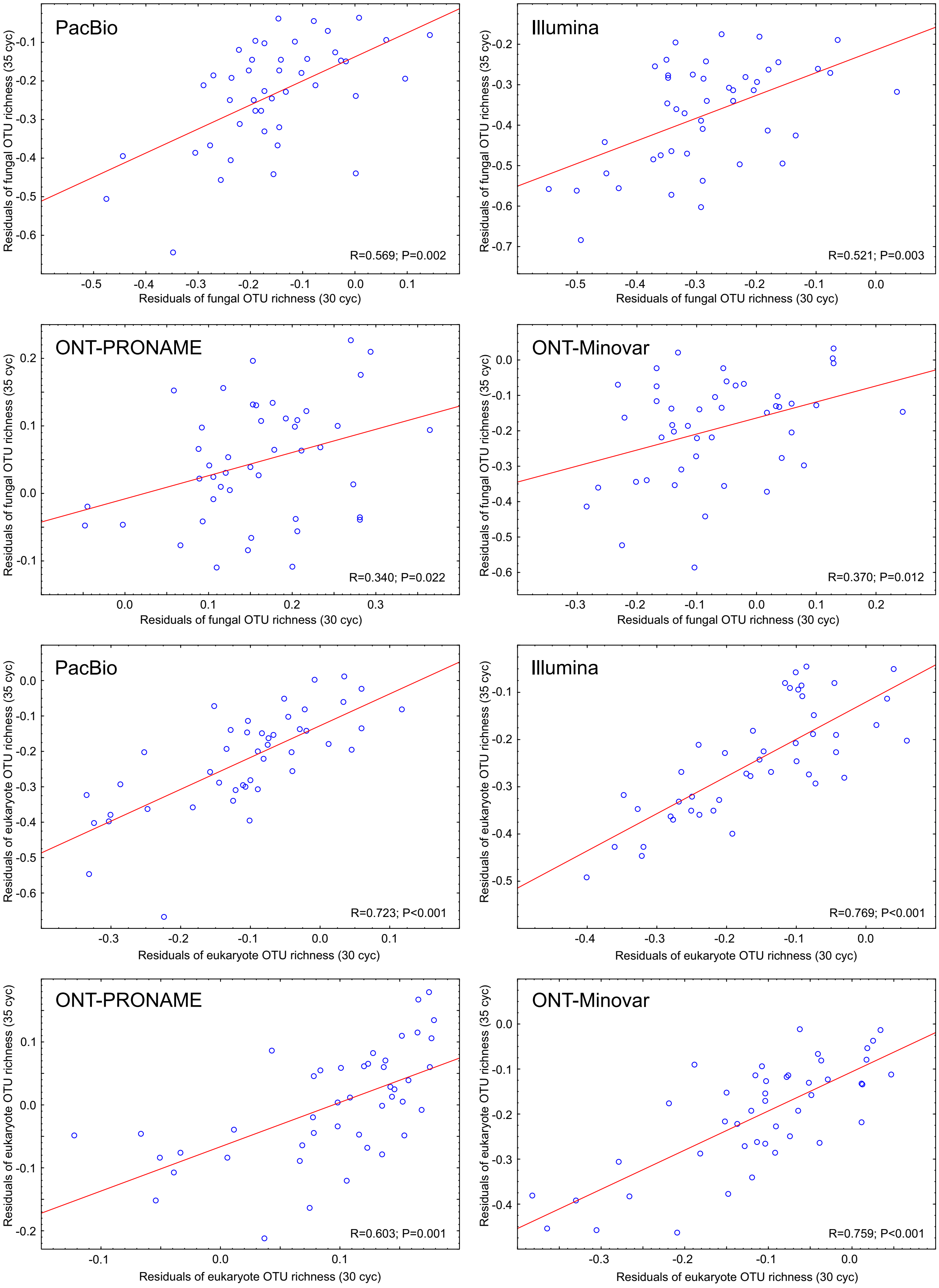


**Figure S3.** Pearson correlations of fungal (top rows) and eukaryote (bottom rows) OTU richness residuals based on 30 and 35 PCR cycles.

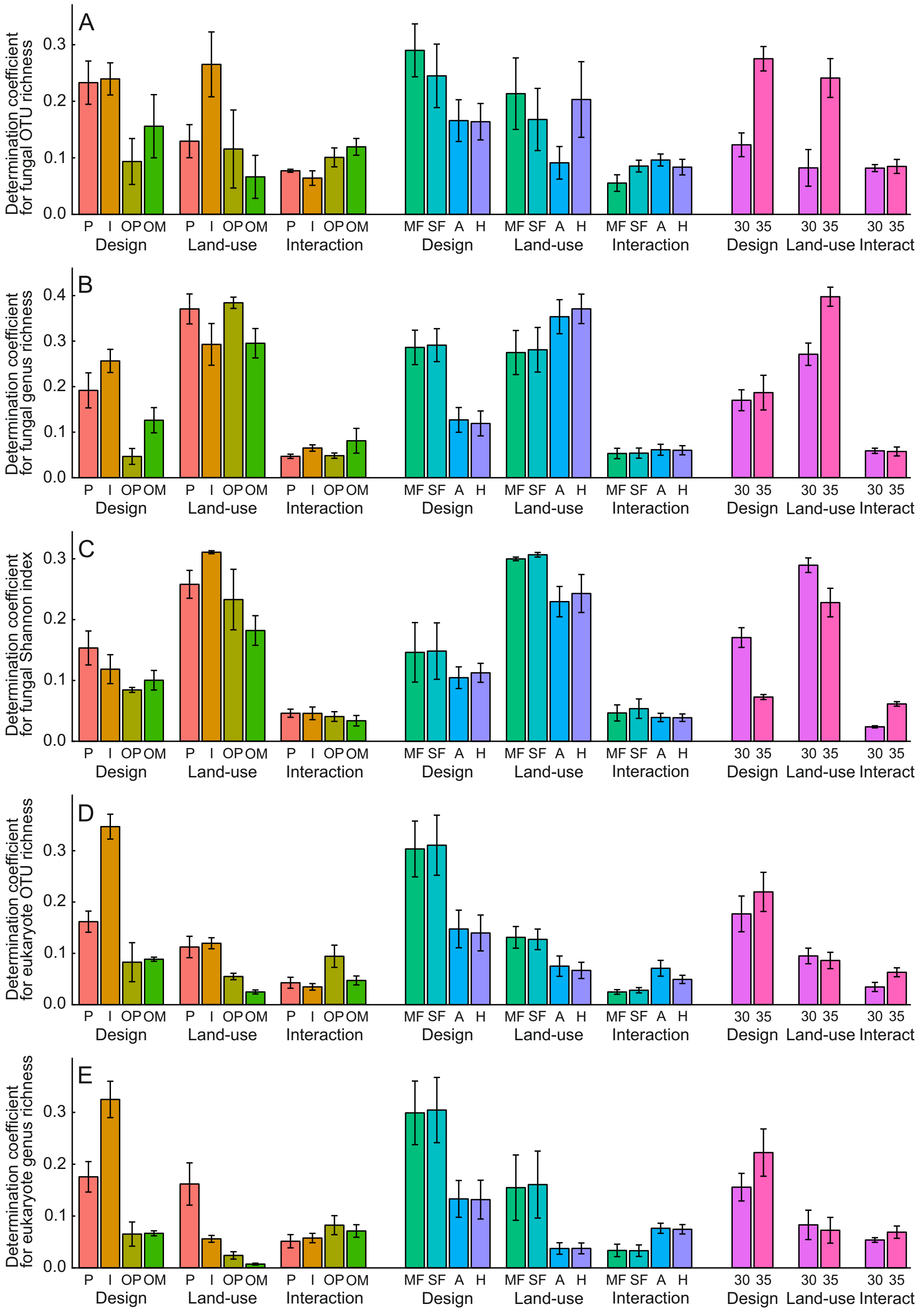


**Figure S4.** Relative effect sizes of sequencing platforms (left panels), bioinformatics pipelines (central panels) and amplicon cycle numbers (right panels) for explaining sampling design, land-use and their interaction effects on A) fungal OTU richness, B) fungal genus richness, C) fungal Shannon index, D) eukaryote OTU richness and E) eukaryote genus richness. The boxes and bars represent means and standard errors. Abbreviations: P, PacBio Revio; I, Illumina MiSeq; OP, Oxford Nanopore PRONAME approach;OM, Oxford Nanopore Minovar approach; MF, minimum-filtering pipeline; SF, standard-filtering pipeline; A, ASV pipeline; H, hybrid approach; 30, 30 PCR cycles; 35, 35 PCR cycles.
